# DNA methylation epigenetically regulates gene expression in *Burkholderia cenocepacia*

**DOI:** 10.1101/2020.02.21.960518

**Authors:** Ian Vandenbussche, Andrea Sass, Marta Pinto-Carbó, Olga Mannweiler, Leo Eberl, Tom Coenye

**Affiliations:** Laboratory of Pharmaceutical Microbiology, Ghent University, Ghent, Belgium; Department of Plant and Microbial Microbiology, University of Zurich, Zurich, Switzerland

## Abstract

Respiratory tract infections by the opportunistic pathogen *Burkholderia cenocepacia* often lead to severe lung damage in cystic fibrosis (CF) patients. New insights in how to tackle these infections might emerge from the field of epigenetics, as DNA methylation has shown to be an important regulator of gene expression. The present study focused on two DNA methyltransferases (MTases) in *B. cenocepacia* strains J2315 and K56-2, and their role in regulating gene expression. *In silico* predicted DNA MTase genes BCAL3494 and BCAM0992 were deleted in both strains, and the phenotypes of the resulting deletion mutants were studied: deletion mutant ΔBCAL3494 showed changes in biofilm structure and cell aggregation, ΔBCAM0992 was less motile. *B. cenocepacia* wild type cultures treated with sinefungin, a known DNA MTase inhibitor, exhibited the same phenotype as DNA MTase deletion mutants. Single-Molecule Real-Time sequencing was used to characterize the methylome of *B. cenocepacia*, including methylation at the origin of replication, and motifs CACAG and GTWWAC were identified as targets of BCAL3494 and BCAM0992, respectively. All genes with methylated motifs in their putative promoter region were identified and qPCR experiments showed an upregulation of several genes, including biofilm and motility related genes, in MTase deletion mutants with unmethylated motifs, explaining the observed phenotypes in these mutants. In summary, our data confirm that DNA methylation plays an important role in regulating the expression of *B. cenocepacia* genes involved in biofilm formation and motility.

**Importance:** CF patients diagnosed with *B. cenocepacia* infections often experience rapid deterioration of lung function, known as *cepacia syndrome. B. cenocepacia* has a large multi-replicon genome and a lot remains to be learned about regulation of gene expression in this organism. From studies in other (model) organisms, it is known that epigenetic changes through DNA methylation play an important role in this regulation. The identification of *B. cenocepacia* genes of which the expression is regulated by DNA methylation and identification of the regulatory systems involved in this methylation are likely to lead to new insights in how to tackle *B. cenocepacia* infections in CF patients.

## Introduction

*Burkholderia cenocepacia*, a member of the *Burkholderia cepacia* complex (Bcc), is an aerobic Gram-negative bacterium that can be isolated from soil and water (1–3). *B. cenocepacia* is also known as an opportunistic pathogen in immunocompromised patients (4–6). Infection of the upper airways in cystic fibrosis (CF) patients often leads to severe illness, typically referred to as *cepacia syndrome* (1,7). CF patients diagnosed with *cepacia syndrome* experience a progressive decrease in lung function, often accompanied by bacteremia and sepsis. If left untreated, *cepacia syndrome* can lead to death within weeks (8,9). The genome of *B. cenocepacia* is complex (with usually three large replicons), with a high GC-content (67%) and large size, comprising approximately 8.06 Mb (10). The species has been classified into different phylogenetic clusters and subdivided into lineages, including the highly transmissible ET-12 lineage that harbors *B. cenocepacia* strains J2315 and K56-2 (11,12).

Epigenetics is the study of heritable changes in gene expression without changes in the actual genomic sequence. In bacterial genomes, epigenetic control is exerted by DNA methyltransferase enzymes (MTases) (13–15). DNA MTases originate from restriction-modification (RM) systems, early defense mechanisms in bacteria with an active interplay between endonucleases and DNA MTases, which cleave foreign DNA but protect the own genome. In addition, discovery of orphan DNA MTases, enzymes without a cognate endonuclease, shows that DNA MTases are not exclusively dependent on the presence of the restriction part to function as regulator of gene expression (16).

DNA MTases interact with specific DNA recognition sites and transfer a CH_3_-group from a methyl donor, mostly S-adenosyl methionine, to a cytosine (C_5_-methyl cytosine or N_4_-methyl cytosine) or adenine (N_6_-methyl adenine) base (17,18). As methylated bases change the binding affinity of DNA binding proteins, methylation at regulatory regions allows bacteria to regulate gene expression at the level of transcription (19,20). While both cytosine and adenine methylation occur in eukaryotic and prokaryotic cells, C_5_-methyl cytosine is the archetypal eukaryotic base methylation signature (16,21). Conversely, in prokaryotes, N_6_-methyl adenine is the most important base modification involved in gene expression regulation (22). In addition to this, studies with DNA MTases *Dam* (deoxyadenosine methyltransferase) and *Dcm* (DNA cytosine MTase) in *Escherichia coli* have demonstrated that, besides having a regulatory function, DNA MTases also take part in crucial cellular processes like DNA replication initiation or methyl-directed mismatch repair (21,23).

Detection of (genome-wide) DNA methylation patterns has been challenging in the past. The use of specific restriction enzymes with affinity for methylated sites, followed by a comparison of the resulting fragment lengths gave a good impression of methylation of the treated DNA at one particular area, but global methylation analysis was until recently difficult at best (21,24). The rise of Next-Generation Sequencing and Single-Molecule Real-Time (SMRT) technologies tremendously improved quality of methylome analyses, but it also made it much more accessible (25,26). SMRT Sequencing uses a sequencing-by-synthesis approach with fluorescently labeled nucleotides. Pulse width, the signal of nucleotide incorporation, and interpulse duration, the time between two incorporations, allow to discriminate between incorporated bases and their methylation status (27).

The purpose of the present study is to understand how DNA methylation regulates gene expression in *B. cenocepacia*. To this end, a genome-wide methylome analysis was carried out, and genes under DNA methylation regulation were identified. Interpretation of the working mechanisms of these regulatory systems, might lead to new insights in how to tackle *B. cenocepacia* infections in CF patients.

## Results

### Identification of *B. cenocepacia* DNA MTases

All predicted DNA MTase genes in the *B. cenocepacia* J2315 genome were identified using REBASE (Table S1). DNA MTase genes BCAL3494 and BCAM0992, widely distributed within the genus *Burkholderia*, were selected for further analysis. Gene BCAL3494, located on the first replicon of *B. cenocepacia*, is a type III methyltransferase that is part of a RM-system, together with a restriction enzyme encoded by the neighbouring gene BCAL3493. Gene BCAM0992 is located on the second replicon and apparently does not have any adjacent genes coding for restriction enzymes. The gene is classified as coding for a type II methyltransferase, i.e. the restriction and modification enzymes act separately and are not dependent on each other (28). To investigate the influence of BCAL3494 and BCAM0992 on bacterial physiology, deletion mutants were constructed (Figure S1). For the other DNA MTase genes in *B. cenocepacia* J2315 identified with REBASE (Table S1), homologues in different *Burkholderia* strains could not be found; these genes were not further investigated in the present study.

### Phenotype of mutant strains

BCAL3494 and BCAM0992 were deleted in two *B. cenocepacia* strains, J2315 and K56-2, and the phenotype of the deletion mutants was investigated in detail. No differences in growth between wild type and mutant strains were observed when cultured in phosphate buffered mineral medium (Figure S2). Microscopic analysis clearly showed a different, more clustered biofilm morphology for both BCAL3494 deletion mutants (ΔBCAL3494) compared to wild type strains, whereas the biofilm structure of the BCAM0992 deletion mutants (ΔBCAM0992) did not differ from wild type (Figure 1A). Cell aggregation in planktonic cultures was investigated using flow cytometry (Figure 1B). The degree of aggregation in the BCAL3494 mutant strains was significantly higher (p-value J2315: 0.049, p-value K56-2: 0.001) than in the corresponding wild type strains. Also, the ability to form a pellicle, a biofilm-like structure at the air-liquid interface, was investigated (Figure 1C). Pellicle formation was clearly increased for both ΔBCAL3494 mutants compared to wild type strains and to ΔBCAM0992 mutants. Complemented mutant strains *c*ΔBCAL3494 and *c*ΔBCAM0992 did not differ significantly from wild type in these experiments (Figure S3).

**FIGURE 1.**
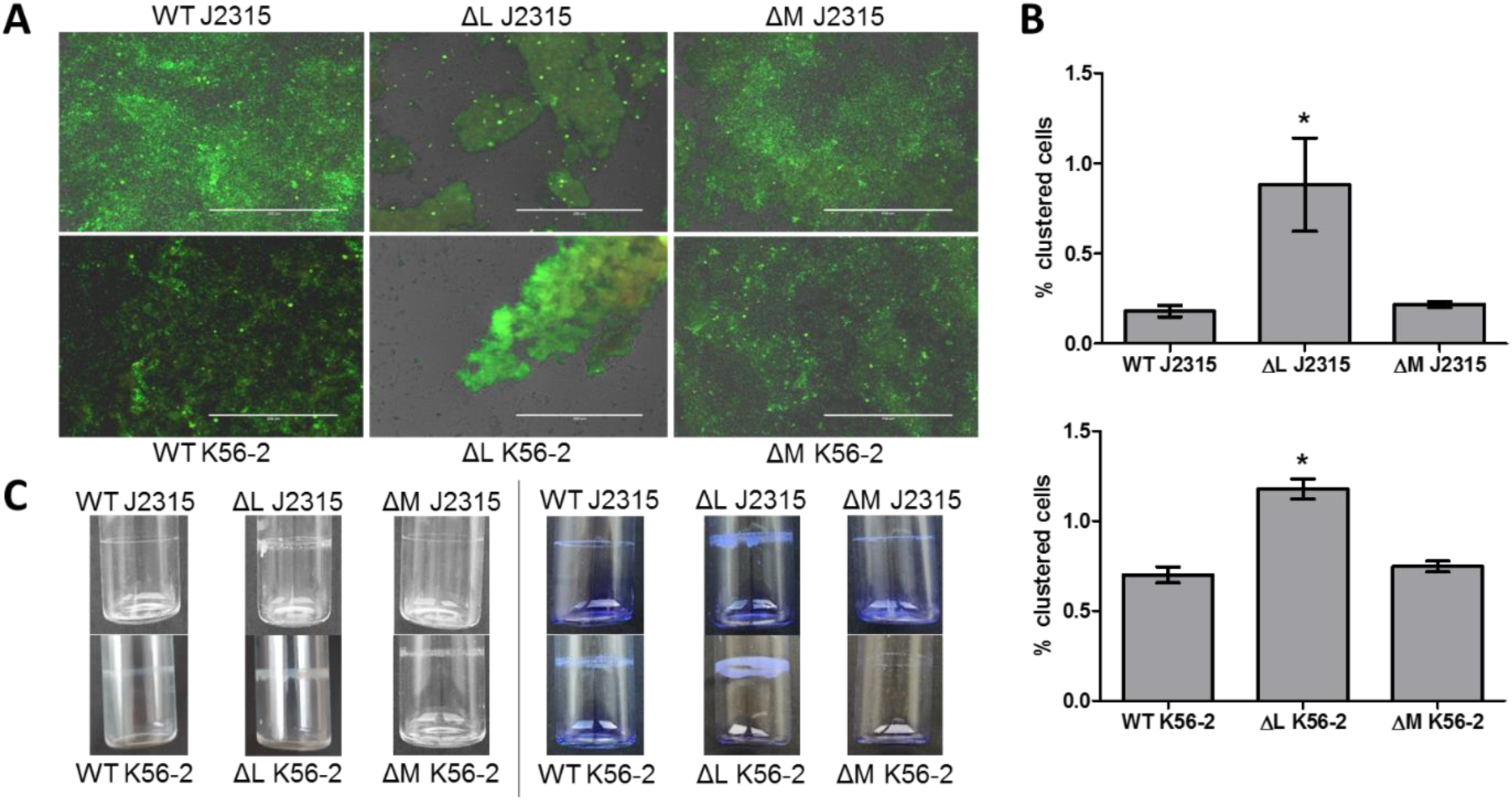
Effect of DNA MTase deletion on biofilm structure, cell aggregation, and pellicle formation in *B. cenocepacia* J2315 and K56-2. (A) Microscopic images of LIVE/DEAD stained biofilms, grown in microtiter plate wells for 24 h. White bar (200 µm) for scale. (B) Clustering of cells in planktonic cultures, quantified with flow cytometry. (C) Pellicle formation inside glass tubes after 24 h of static incubation. Left pictures represent unstained samples, right pictures display pellicles stained with crystal violet. (n=3, * p < 0.05 compared to wild type, error bars represent the standard error of the mean (SEM)). WT: wild type, ΔL: deletion mutant ΔBCAL3494, ΔM: deletion mutant ΔBCAM0992)

Motility of all strains was assessed using a swimming motility assay on 0.3 % agar plates. After 24h (strain K56-2), and 32h (strain J2315), plates were photographed and diameters measured (Figure 2). Diameters were significantly smaller for both ΔBCAM0992 mutants compared to wild type (p-value J2315: 0.002, p-value K56-2 < 0.001). Both ΔBCAL3494 mutants, as well as the complemented mutants, were identical to wild type in this regard. We also investigated swarming motility, but no significant differences between the different strains were observed (Figure S4).

**FIGURE 2.**
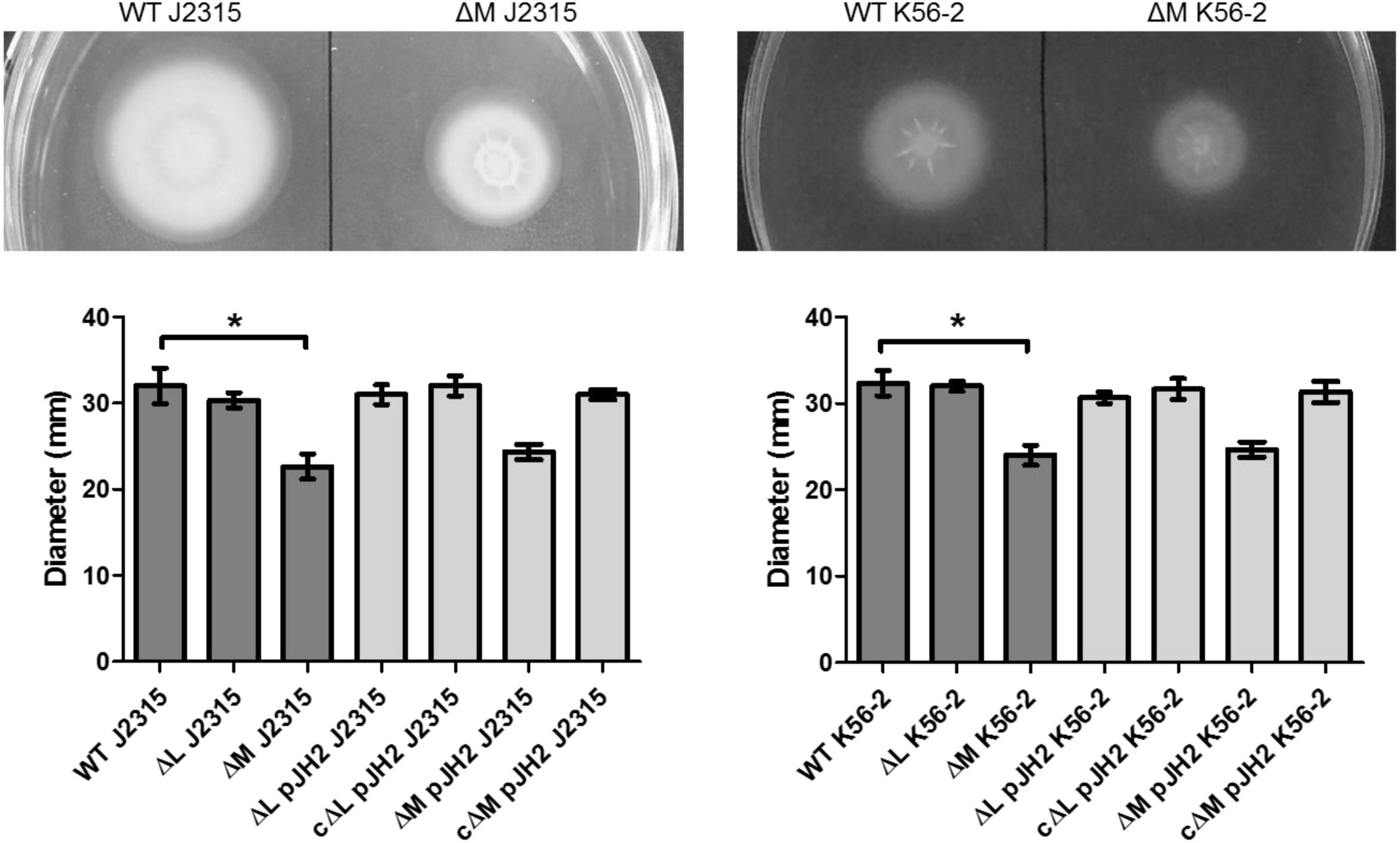
Swimming motility of DNA MTase deletion mutants. Diameters were measured after 24 h (K56-2) or 32 h (J2315). (n=3, * p < 0.05 compared to wild type, error bars represent the SEM. WT: wild type, ΔL: deletion mutant ΔBCAL3494, ΔM: deletion mutant ΔBCAM0992, ΔL pJH2 and ΔM pJH2: mutant strains with empty vector pJH2 (vector control), *c*ΔL pJH2 and *c*ΔM pJH2: deletion mutants complemented with genes BCAL3494 and BCAM0992)

### Effect of the DNA MTase inhibitor sinefungin on methylation-dependent phenotypes

Sinefungin, a structural analog of S-adenosyl methionine and known for blocking base methylation in other bacteria such as *Streptococcus pneumoniae* (29), was used as DNA MTase inhibitor. The minimum inhibitory concentration (MIC) of sinefungin in *B. cenocepacia* J2315 and K56-2 was determined and was found to be higher than 200 µg/mL. Both strains were exposed to sinefungin concentrations below the MIC of sinefungin (50 µg/mL) to assure any effect observed was not due to growth inhibition by sinefungin, and the effect on biofilm formation, pellicle formation, cell aggregation and motility was quantified (Figure 3). Bacteria exposed to sinefungin produced more pellicle mass, showed a higher degree of cell aggregation (p-value: 0.003), had a different biofilm morphology, and were less motile (p-value: 0.004). These findings indicate that chemically blocking DNA methylation or deleting genes responsible for DNA methylation lead to the same phenotypes in *B. cenocepacia* J2315 and K56-2.

**FIGURE 3.**
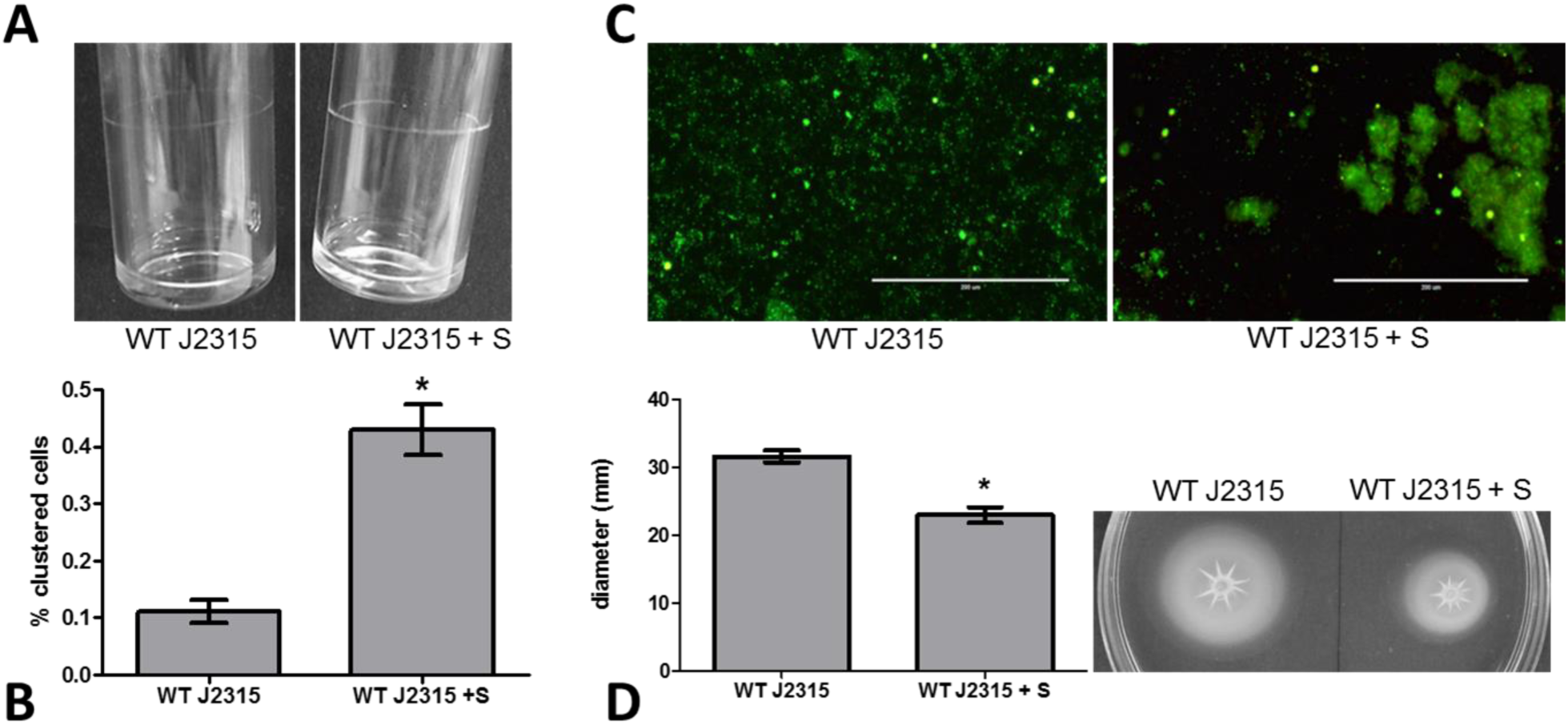
Effect of DNA MTase inhibitor sinefungin on biofilm and pellicle formation, cell aggregation and motility. (A) Pellicle formation inside glass tubes after 24 h of static incubation. (B) Clustering of planktonic cultures analyzed with flow cytometry. (C) Microscopic images of LIVE/DEAD stained biofilms, grown on plastic surfaces in microtiter plates for 24 h. (D) Swimming motility of treated and untreated samples. (n=3, * p < 0.05 compared to wild type, error bars represent the SEM). WT: wild type, +S: medium supplemented with 50 µg/mL sinefungin)

### Methylome analysis

Using SMRT sequencing (PacBio), the complete methylome of *B. cenocepacia* J2315 and K56-2 was identified. Only data for strain J2315 are reported in the following section, as data for strain K56-2 were highly comparable (Figures 4 and 5). Three distinct methylated motifs were identified in the wild type strain: CACAG, GTWWAC, and GCGGCCGC. The CACAG motif was methylated at the fourth position on the forward strand, whereas the GTWWAC motif was methylated at the fifth position on both the forward and reverse strand. Cytosine methylation of the GCGGCCGC motif occurred at the fifth position on both the forward and reverse strand. Although all CACAG and GTWWAC motifs were methylated in the wild type strains, methylation of the CACAG motif was absent in the ΔBCAL3494 deletion mutants, and likewise, no methylation of the GTWWAC motif was seen in the ΔBCAM0992 mutants (Table S2). This demonstrates that MTase BCAL3494 recognizes the CACAG motif, while MTase BCAM0992 recognizes the GTWWAC motif. In contrast, cytosine methylation of the GCGGCCGC motif was observed in wild type and in mutant strains, but the extent of methylation at this motif was highly variable in the four datasets. This suggests that this cytosine methylation occurs randomly, and that the GCGGCCGC motif might present a false positive result of motif analysis due to the high occurrence of short repeats of G and C in the GC-rich *B. cenocepacia* genome. In addition, almost no methylated GCGGCCGC motifs were found in regulatory regions, hinting at only a minor role for cytosine methylation in regulation of gene expression. Therefore, cytosine methylation in *B. cenocepacia* was not further studied.

**FIGURE 4.**
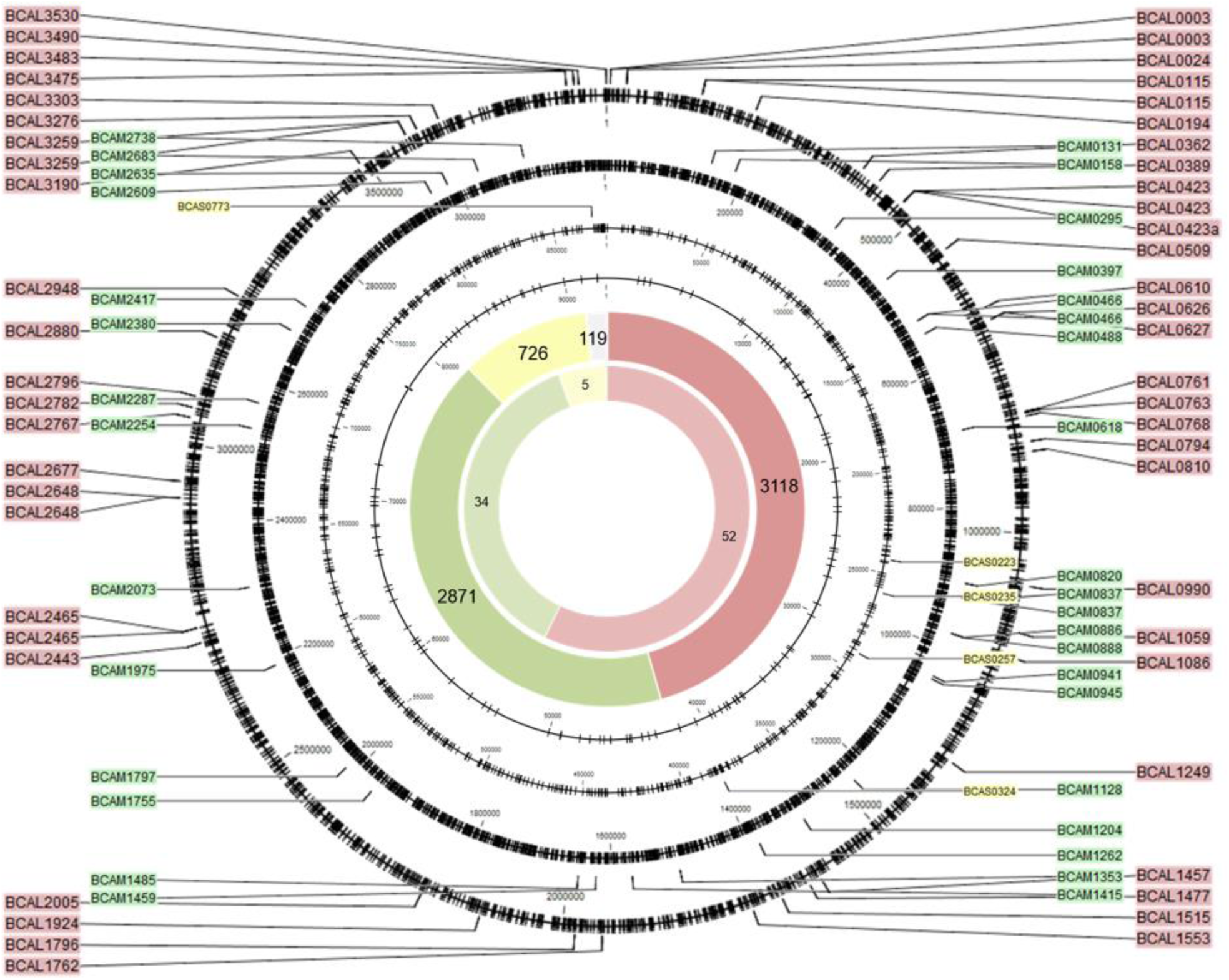
Genomic position of all methylated CACAG motifs. Black circles represent the four replicons of *B. cenocepacia*, black ticks mark the motif locations. The total number of methylated CACAG motifs and methylated CACAG motifs in promoter regions, per replicon (red: replicon 1, green: replicon 2, yellow: replicon 3, and grey: plasmid), is shown on the large and small inner circle, respectively. The position and names of genes with methylated promoter regions is indicated with colored labels (same color code).

**FIGURE 5.**
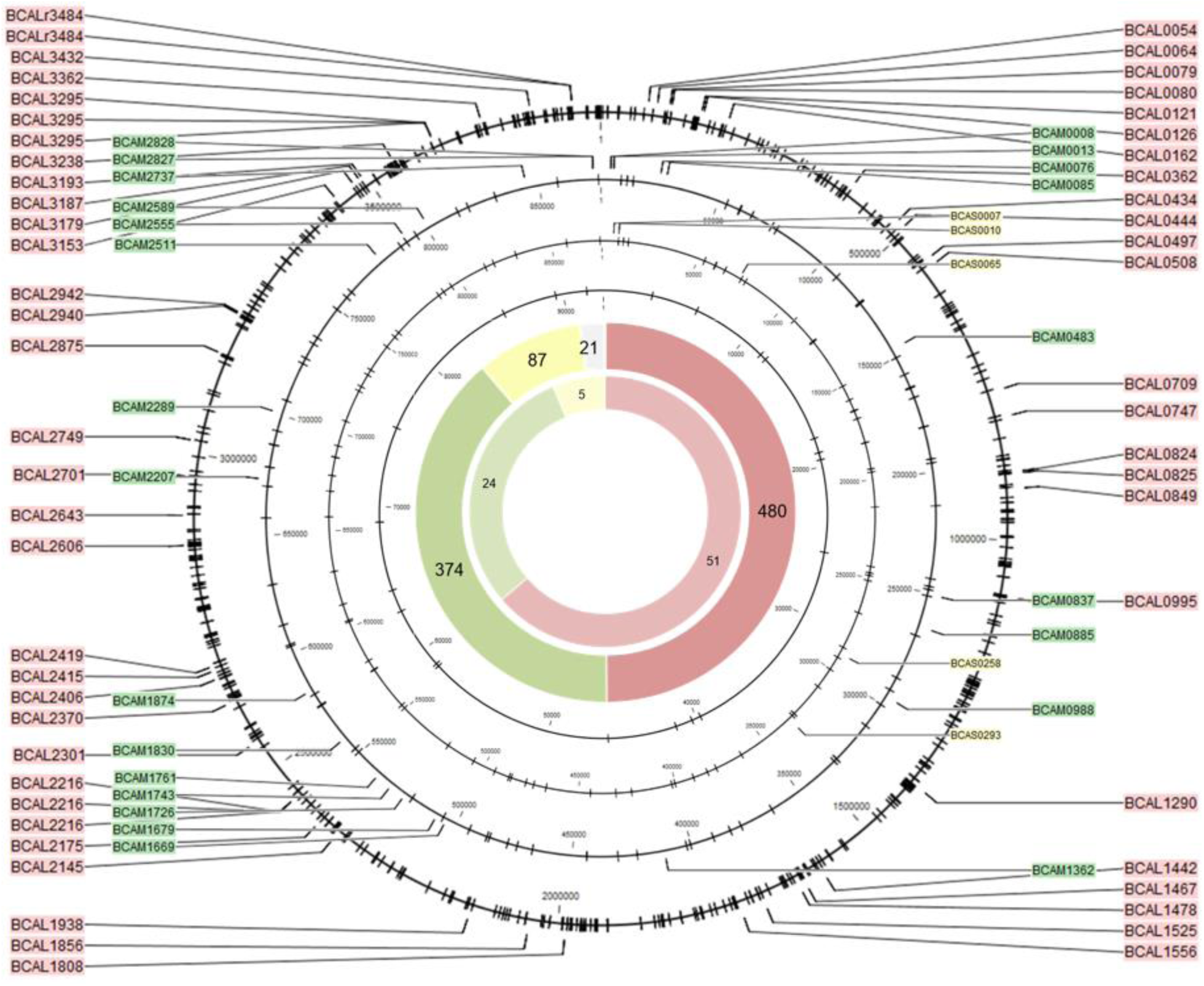
Genomic position of all methylated GTWWAC motifs. Black circles represent the four replicons of *B. cenocepacia*, black ticks mark the motif locations. The total number of methylated GTWWAC motifs and methylated GTWWAC motifs in promoter regions, per replicon (red: replicon 1, green: replicon 2, yellow: replicon 3, and grey: plasmid), is shown on the large and small inner circle, respectively. The position and names of genes with methylated promoter regions is indicated with colored labels (same color code).

The location of every methylated CACAG and GTWWAC motif was mapped (Figures 4 and 5). A total of 6834 methylated CACAG motifs and 961 methylated GTWWAC motifs was found, of which the majority was present on the first replicon (CACAG: 45.6 %, GTWWAC: 49.9 %), followed by the second replicon (CACAG: 42.1 %, GTWWAC: 38.9 %), the third replicon (CACAG: 10.6 %, GTWWAC: 9.0 %), and the plasmid (CACAG: 1.7 %, GTWWAC: 2.2 %). Subsequently, all genes with methylated motifs in their promoter region, here defined as 60 bases upstream of the transcription start site, were identified. 91 promoter regions contained methylated CACAG motifs, and 80 promoter regions contained methylated GTWWAC motifs, with most of the motifs being present on the first replicon (Figures 4 and 5). Functional classes of genes found in the dataset of genes with methylated promoter include genes involved in intermediary metabolism, regulation, and transport (Tables S3 and S4).

Virtual Footprint was used to elucidate to which transcription factor (TF) binding sites the discovered methylation motifs CACAG and GTWWAC showed any similarity. Data output of the analysis is listed in Table S5. Sequences that contain methylation motif CACAG were similar to the binding site of *E. coli* K12 GlpR, while GTWWAC-containing sequences were similar to binding sites of several other *E. coli* K12 TFs, including ArcA, OxyR, Fis and Fur.

### Expression of genes with a methylated promoter

The expression level of genes with methylated promoter regions was determined in wild type and mutant strains, using qPCR. Expression data of genes with methylated promoter regions are listed in Tables 1 and 2. Volcano plots (Figure S5, fold changes plotted against corresponding p-values) show that most genes tested were upregulated in the mutants compared to the wild type strains. Six of these genes were significantly upregulated in mutants of both strain backgrounds: BCAL1515, BCAL2465, and BCAM0820 were upregulated in ΔBCAL3494, whereas genes BCAL0079, BCAL2415, and BCAM1362 were upregulated in ΔBCAM0992. Four additional genes were upregulated in K56-2 mutants only: BCAL0423, BCAM2738, and BCAS0223 were upregulated in ΔBCAL3494, BCAL1556 in ΔBCAM0992. Subsequently, the methylated promoter regions of these genes were analyzed in detail (Figure 6). In most cases, the methylated motif was in close proximity of the −10 or −30/35 element in bacterial promoter regions.

**TABLE 1.**
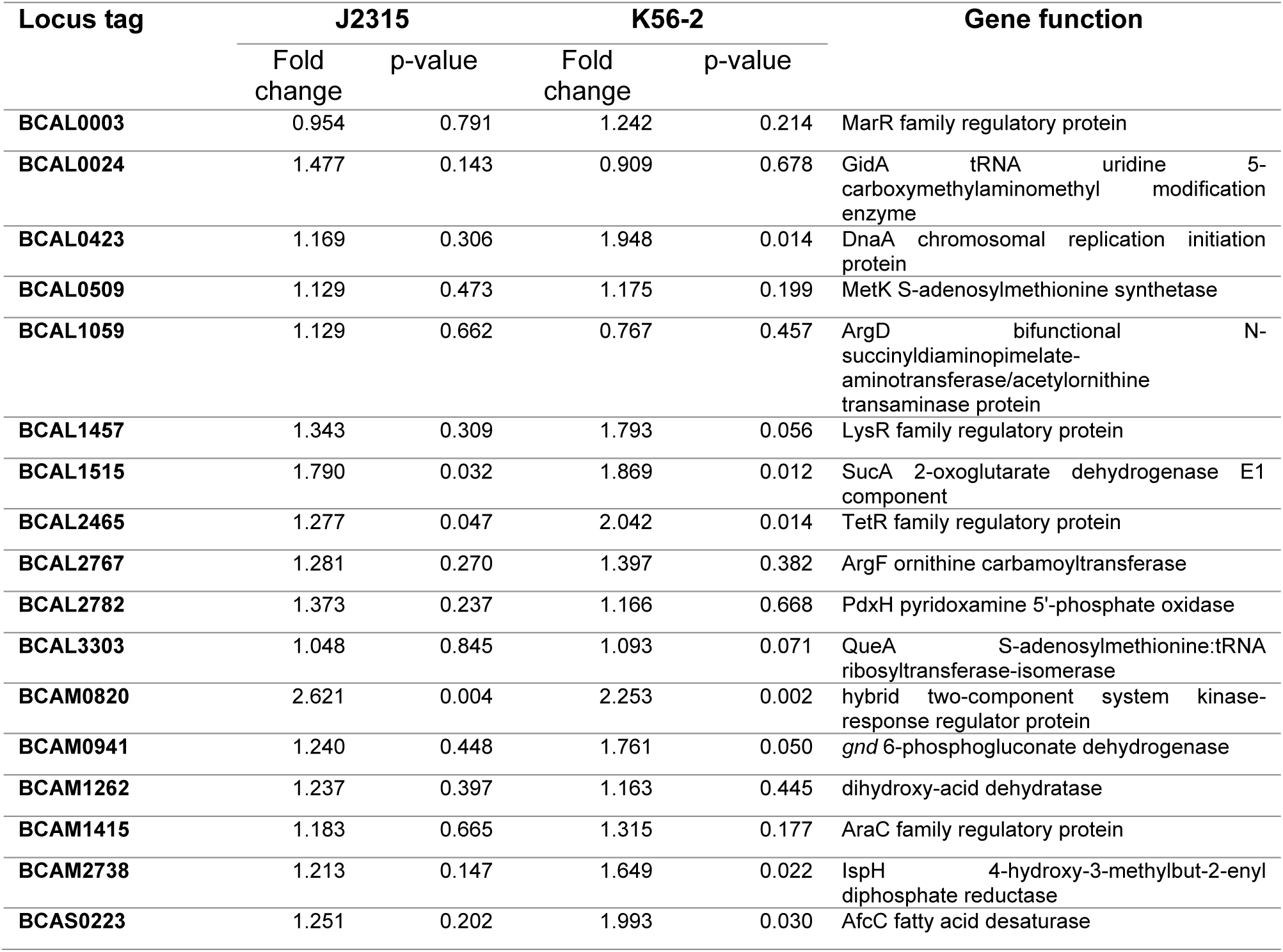
Expression changes of genes with a methylated CACAG motif in their promoter region in deletion mutants compared to wild type.

**TABLE 2.** Expression changes of genes with a methylated GTWWAC motif in their promoter region in deletion mutants compared to wild type.

| Locus tag | J2315 |  | K56-2 |  | Gene function |
| --- | --- | --- | --- | --- | --- |
|  | Fold change | p-value | Fold change | p-value |  |
| BCAL0054 | 0.487 | 0.127 | 0.641 | 0.107 | MerR family regulatory protein |
| BCAL0079 | 2.838 | 0.020 | 3.074 | 0.005 | *rep* ATP-dependent DNA helicase |
| BCAL0126 | 0.845 | 0.356 | 0.801 | 0.607 | MotA chemotaxis protein |
| BCAL0162 | 0.139 | 0.634 | 0.133 | 0.478 | GmhA phosphoheptose isomerase |
| BCAL0508 | 1.015 | 0.934 | 1.721 | 0.137 | LpxL lipid A biosynthesis myristoyl acyltransferase |
| BCAL0709 | 1.599 | 0.104 | 0.763 | 0.430 | LipB lipoate-protein ligase B |
| BCAL1556 | 1.611 | 0.171 | 1.690 | 0.006 | RpiA ribose-5-phosphate isomerase A |
| BCAL2406 | 1.693 | 0.273 | 1.009 | 0.938 | WabR putative glycosyltransferase |
| BCAL2415 | 2.819 | 0.006 | 6.029 | 0.001 | PurT phosphoribosylglycinamide formyltransferase 2 |
| BCAL2701 | 0.613 | 0.076 | 1.519 | 0.170 | ArgD acetylornithine transaminase protein |
| BCAL2942 | 1.143 | 0.474 | 1.451 | 0.274 | CysM cysteine synthase B |
| BCAM0076 | 1.630 | 0.112 | 1.358 | 0.051 | TetR family regulatory protein |
| BCAM1362 | 1.959 | 0.025 | 1.516 | 0.004 | putative penicillin-binding protein |
| BCAS0258 | 1.247 | 0.451 | 1.141 | 0.438 | GntR family regulatory protein |

**FIGURE 6.**
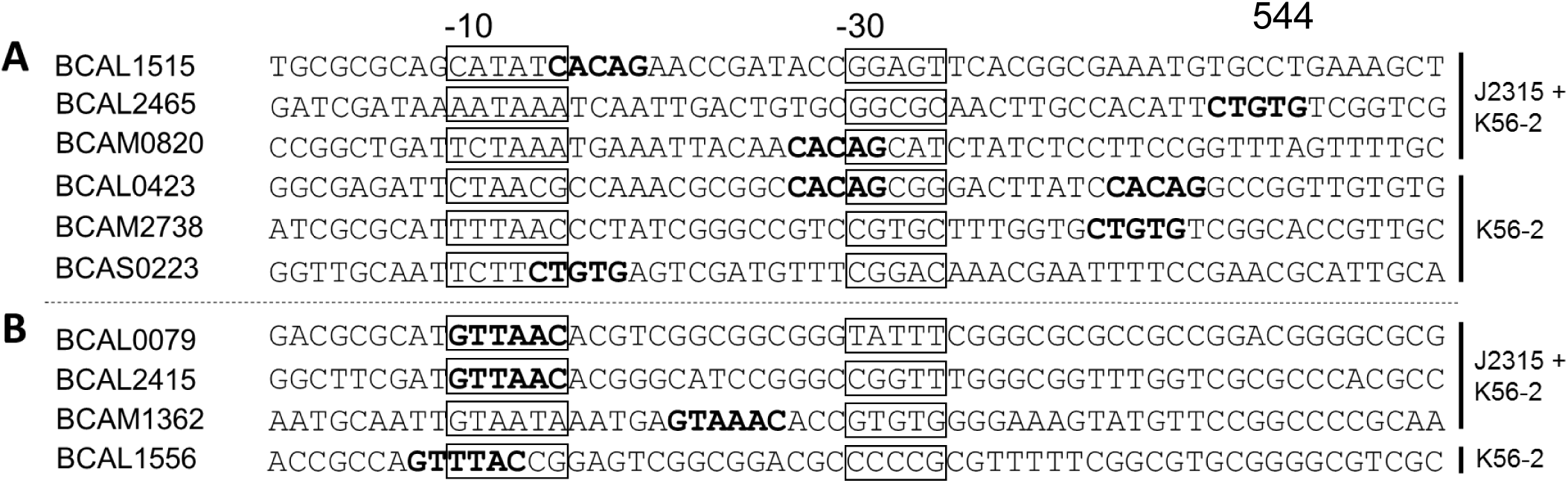
Position of methylated motifs relative to gene start for genes of which the expression is upregulated in DNA MTase deletion mutants. (A) Genes with methylated CACAG motifs in their corresponding promoter region. (B) Genes with methylated GTWWAC motifs in their corresponding promoter region. The motifs are marked in bold, the position of −10 and −30/35 elements in bacterial promoters are framed. (‘J2315 + K56-2’: upregulation in both strains, ‘K56-2’: upregulation in strain K56-2 only)

To confirm that the presence of methylation close to the −10 or −30/35 element influences transcription and therefore gene expression in *B. cenocepacia*, translational eGFP reporter fusions were constructed and eGFP production was quantified. The eGFP production in strains harboring different plasmids is shown in Figure 7. As expected, the production of eGFP, driven by the promoters of genes BCAL1515, BCAM0820, and BCAL0079, was significantly (p = 0.001, p = 0.014, p = 0.002, respectively) increased in the deletion mutant for which an upregulation of these genes was observed using qPCR experiments (Figure 7).

**FIGURE 7.**
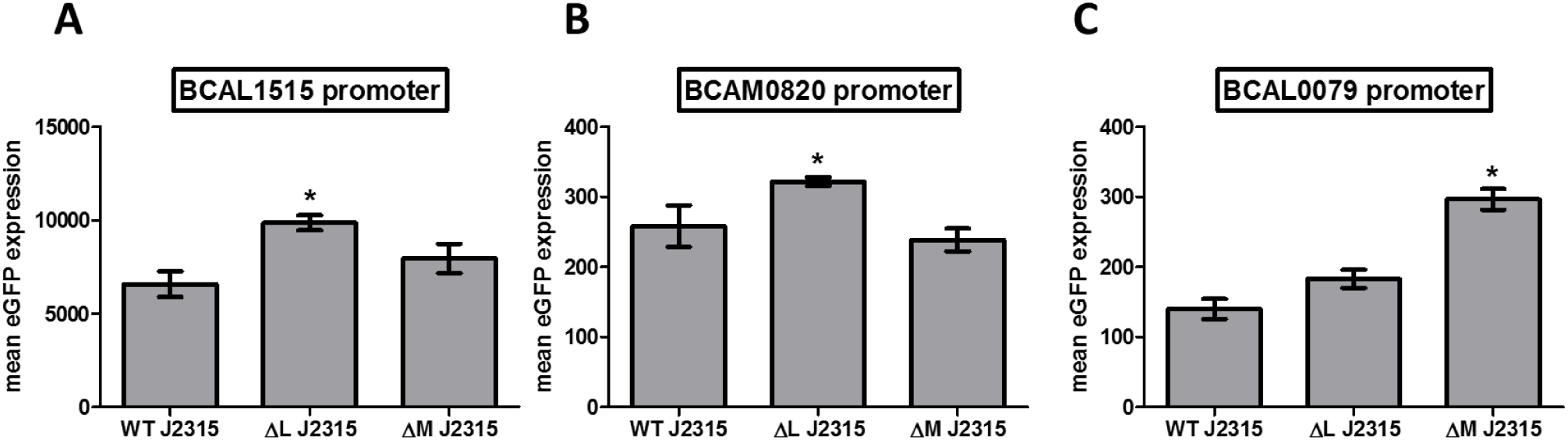
eGFP production in *B. cenocepacia* J2315 strains harboring a pJH2 plasmid that contains a BCAL1515 promoter-eGFP construct (A), a BCAM0820 promoter-eGFP construct (B), or a BCAL0079 promoter-eGFP construct (C). BCAL 1515 and BCAM0820 are associated with methylation of the CACAG motif by DNA MTase BCAL3494, BCAL0079 is associated with methylation of the GTWWAC motif by DNA MTase BCAL0992 (n=3, * p < 0.05 compared to wild type, error bars represent the SEM). WT: wild type, ΔL: deletion mutant ΔBCAL3494, ΔM: deletion mutant ΔBCAM0992).

### DNA methylation in the origin of replication

DNA methylation was detected in all origins of replication of *B. cenocepacia* (Figure 8). Similar methylation patterns were observed in the origins of the different replicons. A previously discovered 7-mer (CTGTGCA) that can be found in all replication origins (30), contains a CACAG methylation motif on the antisense strand. This motif was also found at the 3’-end of almost every DnaA box. These boxes are bound by DnaA proteins, essential for DNA unwinding and chromosome replication initiation (31). Also, the GTWWAC motif was found in proximity of the replication origins, consequently the origins in *B. cenocepacia* represent methylation-rich regions. Whereas methylated CACAG motifs were found throughout the origins of replication, the position of the GTWWAC methylation was unique in all replicons and at least two GTWWAC motifs were found in between two CACAG methylated DnaA boxes. In contrast to the origins of the three larger replicons, the origin of replication of the plasmid contained only one CACAG methylated DnaA box.

**FIGURE 8.**
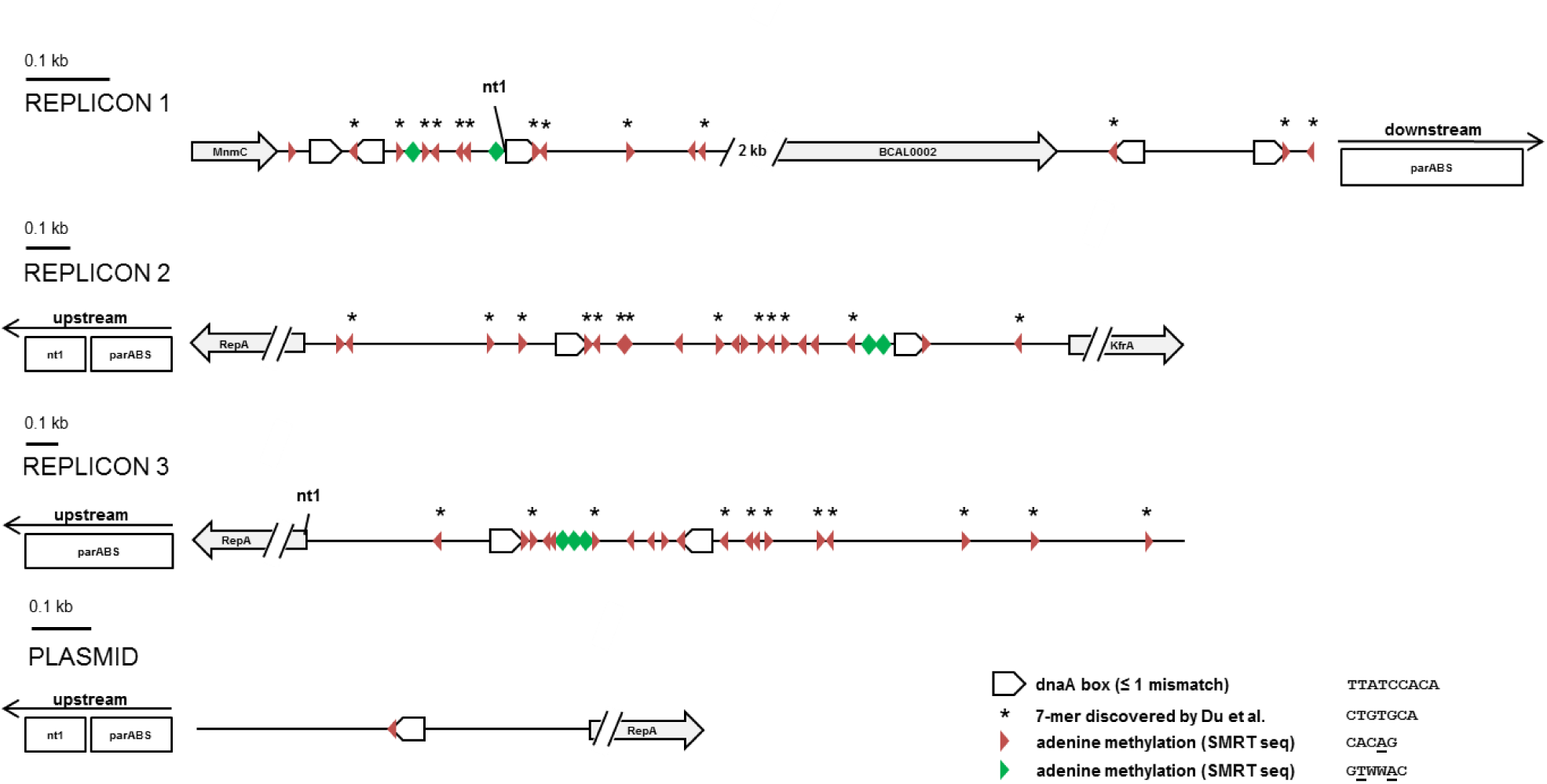
Methylation in the origin of replication of the different replicons in *B. cenocepacia* J2315. SMRT sequencing was used to detect methylated CACAG (red triangles) and GTWWAC (green triangles) motifs within these regions. DnaA boxes (TTATCCACA, consensus sequence of DnaA boxes in *E. coli*) are indicated in the figure. CACAG motifs were frequently found to be part of a previously discovered 7-mer (sense: CTGTGCA, antisense: TGCACAG) (30). The positions of these 7-mers are indicated with an asterisk. (nt1, nucleotide 1; parABS genes, responsible for chromosome segregation in *B. cenocepacia*)

## Discussion

Despite the growing knowledge of DNA methylation in prokaryotes (15), the role of DNA MTases in regulating gene expression in *B. cenocepacia* remains to be revealed. In the present study, we identified two DNA MTases (BCAL3494 and BCAM0992), and mutants in which these genes were deleted, showed differences in biofilm formation and motility. In addition, when methylation was blocked by the DNA MTase inhibitor sinefungin (32), the same phenotypic differences were observed. These findings demonstrate that epigenetic control of gene expression by MTases play an important role in controlling certain phenotypes. Similar results have been reported in *Salmonella enterica*, where DNA methylation is crucial for optimal pellicle and biofilm production (33).

Methylome analysis showed that mutants in which MTase ΔBCAL3494 or ΔBCAM0992 were inactivated, lacked adenine methylation in specific motifs. MTase BCAL3494 was specifically linked to methylation of the CACAG motif, MTase BCAM0992 to methylation of the GTWWAC motif. This strategy of DNA methylation analysis, in which the methylome of strains lacking MTases is determined, has been used in various bacteria, as it is an effective way to find associations between predicted MTases and genome-wide methylation motifs (34,35). For example, several methylation motifs were identified in *Burkholderia pseudomallei*, including motifs CACAG and GTWWAC (36). Two of the *B. pseudomallei* MTases (M.BpsI and M.BpsII) are homologous to the *B. cenocepacia* MTases BCAL3494 and BCAM0992. In *Ralstonia solanacearum*, an important plant pathogen that is phylogenetically related to *B. cenocepacia*, the GTWWAC methylation motif co-occurs with the respective homolog of the BCAM0992 MTase, whereas a BCAL3494 MTase homolog and methylation of CACAG are absent (37). As in *B. cenocepacia*, the BCAM0992 homolog in *R. solanacearum* is an orphan DNA MTase. Analysis of cytosine methylation suggests that cytosine is more likely to be methylated at random instead of at specific motifs, and is likely not having a major regulatory function. Also, GC-rich genomes complicate the search for specific cytosine motifs.

Previous epigenetic research demonstrated that there is a negative correlation between methylation in promoters and transcription (38). To uncover the role of DNA methylation in regulation of *B. cenocepacia* gene expression, all methylated motifs in promoter regions were identified. The data obtained in the present study indicates that gene expression was upregulated in DNA MTase mutants, suggesting that adenine DNA methylation in *B. cenocepacia* affects gene expression by a mechanism inhibiting transcription. In both prokaryotes and eukaryotes, adenine and cytosine methylation are involved in blocking (or enhancing) the binding of RNA polymerase to DNA (15,21,39), and especially methylation near the −10 and −30/35 elements in the promoter region seems to be important for affecting RNA polymerase binding (40). We found that, also in *B. cenocepacia*, methylated motifs (CACAG and GTWWAC) are found close to, or in these elements.

BCAM0820, upregulated in the J2315 and K56-2 ΔBCAL3494 mutant, is a two-component response regulator, the first gene of an operon homologous to the Wsp chemosensory system involved in biofilm formation in *Pseudomonas aeruginosa* (41). BCAM0820 is homologous to WspR, but lacks the diguanylate cyclase domain. During an experimental evolution study in which *B. cenocepacia* HI2424 biofilms were grown on beads, mutations within the *wsp* gene cluster occurred in different clones; these were associated with increased pellicle formation and increased biofilm formation on beads. This demonstrates that the Wsp cluster is involved in pellicle formation in *B. cenocepacia* (42,43), and the upregulation of BCAM0820 could explain the differences in pellicle and biofilm formation between the wild type strains and the ΔBCAL3494 deletion mutants observed in the present study. Interestingly, BCAL1515, encoding 2-oxoglutarate dehydrogenase (SucA) and upregulated in ΔBCAL3494, also acquired mutations in the course of the experimental evolution study (43), but the role of this gene in biofilm formation has not been further explored. BCAL0079, upregulated in the ΔBCAM0992 mutants, is annotated as a DNA helicase gene (*rep*). Besides unwinding DNA during DNA replication, Rep plays a role in swimming motility in *E. coli* (44). The reduced motility observed in the ΔBCAM0992 mutants suggests that Rep may also affect motility in *B. cenocepacia*, although this remains to be confirmed.

Measurement of eGFP production in translational fusion mutants revealed that mutants with constructs containing the BCAL1515, BCAM0820, or BCAL0079 promoter, showed a significant increase in eGFP production compared to wild type, thereby supporting our hypothesis of gene expression regulation by DNA methylation. *In silico* analyses predict that sequences containing methylation motifs are similar to binding sites of TF in *E. coli* K12, and it is plausible that these sequences are also part of TF binding sites in *B. cenocepacia*, allowing us to propose a possible mechanism of gene expression regulation (Figure 9). TFs that bind close to the −10 and −35 region often act as transcriptional repressors (45). Therefore, a methylated promoter region could promote binding of a repressor (46), and sterically hinder RNA polymerase (OFF state), whereas an absence of methylation would allow binding of the initiation factor sigma to the promoter, which in turn could lead to binding of RNA polymerase and initiation of transcription (ON state).

**FIGURE 9.**
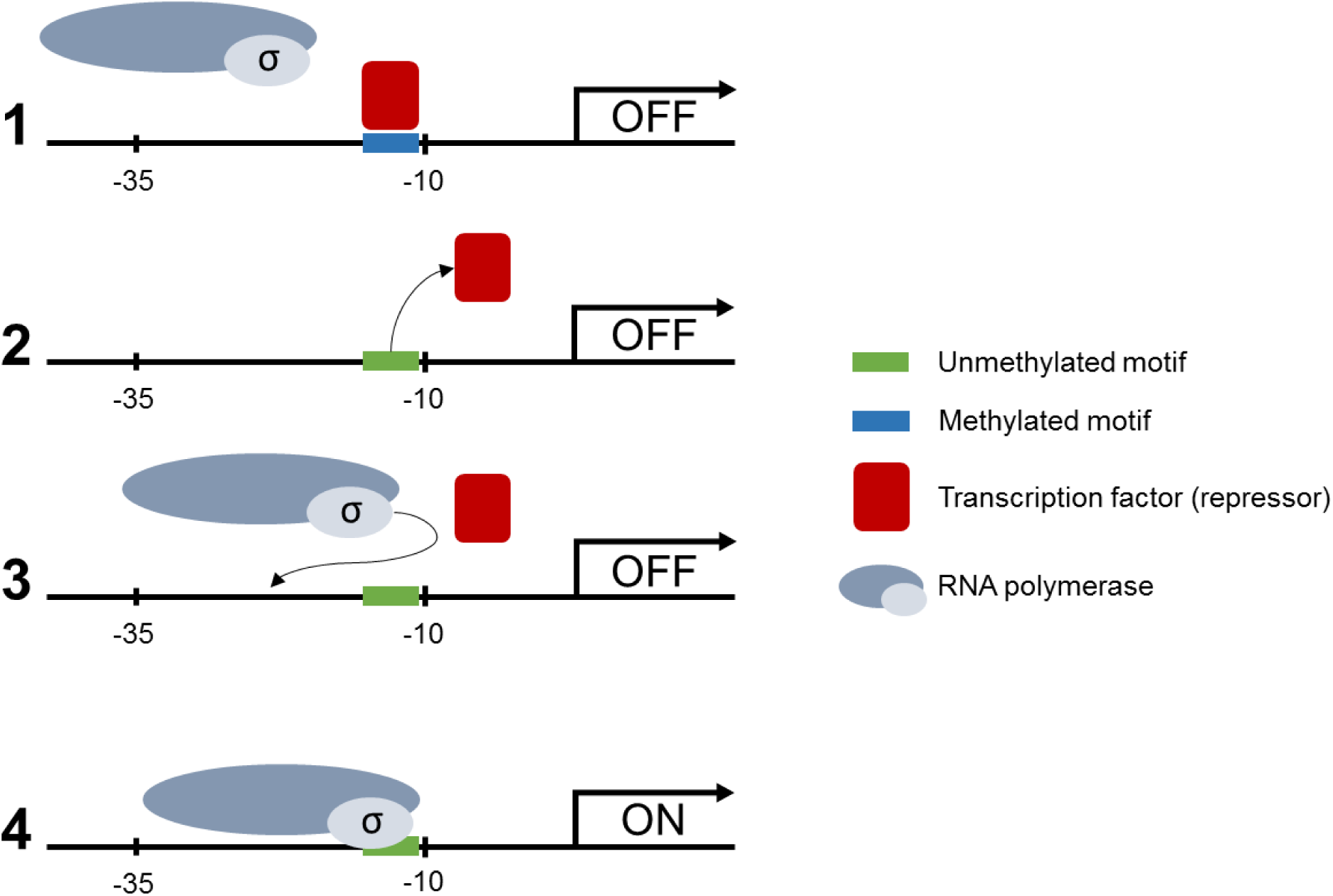
Proposed mechanism of regulation of gene expression in *B. cenocepacia*. (1) Methylated motifs in the promoter region of the gene are bound by a TF, acting as repressor (OFF state). (2) In absence of methylation in the promoter region, the TF dissociates from the motif and vacates the promoter region. (3) The sigma factor is no longer sterically hindered by a repressor and is able to bind to the promoter region. (4) RNA polymerase can access the promoter region and start transcription of the gene (ON state).

The role of DNA methylation in prokaryotes in multifaceted. Besides gene expression regulation and a role in DNA mismatch repair in Gram-positive bacteria (47), DNA methylation has also been implicated in the coordination of replication initiation. Results of the present study seem to confirm this, as the *rep* gene, necessary for replication, was found to be under epigenetic control by DNA methylation. In *E. coli*, GATC motifs, omnipresent in the replication origin, are prone to adenine methylation. The motifs are found within DnaA boxes, essential for binding of the DnaA protein and initiation of replication. The methylation state of each of these GATC motifs changes the affinity of DnaA and sequestering-protein SeqA for the DnaA box. Immediately after replication, GATC motifs are hemi-methylated, which leads to sequestration of the DnaA boxes by SeqA and prevents the process of replication to be reinitiated (21). The occurrence of methylated motifs in the vicinity of the origins of replication of the four replicons in *B. cenocepacia* was studied to check for a link between DNA methylation and coordination of the replication process. An enrichment of the CACAG motif was observed in the origin of replication of all replicons. The motif was found to be part of a bigger sequence that has previously been reported as a recurring 7-mer (30), without known function. In addition, the origin of replication of the different replicons showed high similarities in methylation patterns, raising the possibility of replication coordination by DNA methylation.

In conclusion, we have demonstrated that DNA methylation plays a role in regulation of gene expression in *B. cenocepacia*. DNA MTases BCAL3494 and BCAM0992 are essential for methylation of the *B. cenocepacia* genome, and are responsible for methylation of base motifs CACAG and GTWWAC, respectively. In absence of methylation, expression of certain genes in affected and this results in altered phenotypes (including cell clustering, biofilm formation, and motility). Finally, recurrent methylation patterns were detected in all origins of replication, which suggests an additional role of DNA methylation in replication regulation.

## Materials and Methods

### Strains and culture conditions

All strains and plasmids used in this study are listed in Table 3. *B. cenocepacia* strains were cultivated in phosphate buffered mineral medium (2.00 g/L NH_4_Cl, 4.25 g/L K_2_HPO_4_.3H_2_O (ChemLab), 1.00 g/L NaH_2_PO_4_.H_2_O, 0.10 g/L nitriloacetic acid, 0.0030 g/L MnSO_4_.H_2_O, 0.0030 g/L ZnSO_4_.7H_2_O, 0.0010 g/L CoSO_4_.7H_2_O, 0.20 g/L MgSO_4_.7H_2_O, 0.012 g/L FeSO_4_.7H_2_O (Sigma-Aldrich), 5 g/L Yeast Extract (Lab M), 2 g/L Casamino Acids (BD Biosciences), and 5 g/L glycerol (Scharlab)). LB medium (Luria Bertani medium with 5g/L NaCl, Sigma-Aldrich) was used for maintenance of *E. coli* strains and during specific stages of the gene deletion procedure (see below) where antibiotic selection with tetracycline (250 µg/mL) (Sigma-Aldrich) was desired. Prior to phenotypic experiments, liquid overnight (ON) cultures were grown in a shaker incubator (100 rpm) at 37 °C.

**TABLE 3.** Bacterial strains and plasmids.

| Strain/Plasmid | Description | Abbreviation | Source |
| --- | --- | --- | --- |
| ***B. cenocepacia*** |  |  |  |
| J2315 | CF sputum isolate | WT J2315 | LMG16656 |
| J2315 ΔBCAL3494 | BCAL3494 MTase deletion mutant | ΔL J2315 | this study |
| J2315 ΔBCAM0992 | BCAM0992 MTase deletion mutant | ΔM J2315 | this study |
| J2315 ΔBCAL3494 pJH2 | BCAL3494 MTase mutant with empty pJH2 vector | ΔL pJH2 J2315 | this study |
| J2315 ΔBCAL3494 pJH2 + BCAL3494 | BCAL3494 MTase complemented deletion mutant | cΔL pJH2 J2315 | this study |
| J2315 ΔBCAM0992 pJH2 | BCAM0992 MTase mutant with empty pJH2 vector | ΔM pJH2 J2315 | this study |
| J2315 ΔBCAM0992 pJH2 + BCAM0992 | BCAM0992 MTase complemented deletion mutant | cΔM pJH2 J2315 | this study |
| K56-2 | CF sputum isolate | WT K56-2 | LMG 18863 |
| K56-2 ΔBCAL3494 | BCAL3494 MTase deletion mutant | ΔL K56-2 | this study |
| K56-2 ΔBCAM0992 | BCAM0992 MTase deletion mutant | ΔM K56-2 | this study |
| K56-2 ΔBCAL3494 pJH2 | BCAL3494 MTase mutant with empty pJH2 vector | ΔL pJH2 K56-2 | this study |
| K56-2 ΔBCAL3494 pJH2 + BCAL3494 | BCAL3494 MTase complemented deletion mutant | cΔL pJH2 K56-2 | this study |
| K56-2 ΔBCAM0992 pJH2 | BCAM0992 MTase mutant with empty pJH2 vector | ΔM pJH2 K56-2 | this study |
| K56-2 ΔBCAM0992 pJH2 + BCAM0992 | BCAM0992 MTase complemented deletion mutant | cΔM pJH2 K56-2 | this study |
| ***E. coli*** |  |  |  |
| DH5α | Maintenance of replicative plasmids | | lab stock |
| One Shot PIR2 | Maintenance of suicide plasmids with ori<sub>R6K</sub> | PIR2 | ThermoFisher |
| **Plasmids** |  |  |  |
| pGPI-Scel-XCm | Suicide plasmid, Tp<sup>r</sup>, Cm<sup>r</sup>, I-SceI restriction site, ori<sub>R6K</sub> | pGPI | 44,45 |
| pDAI-Scel-SacB | Replicative plasmid, Tet<sup>r</sup>, I-SceI nuclease, counterselectable marker SacB, ori<sub>pBBR1</sub> | pDAI | 44,45 |
| pRK2013 | Helper plasmid, Km<sup>r</sup>, ori<sub>colEI</sub> | pRK | 44,45 |
| pJH2 | Broad-range translational fusion vector, Cm<sup>r</sup>, fluorescent marker eGFP: complementation of ΔBCAL3494 | | 46 |
| pSCrhaB2 | Broad-range translational fusion vector, Tp<sup>r</sup>, rhaR, rhaS-P<sub>rhaB</sub>, ori<sub>pBBR1</sub>: complementation of ΔBCAM0992 | | 47 |
| pGPI + BCAL3494 upstream sequence | pGPI-Scel-XCm with ligated upstream sequence BCAL3494, used during deletion | pGPI<sub>UL</sub> | this study |
| pGPI + BCAL3494 upstream and downstream sequence | pGPI-Scel-XCm with ligated upstream and downstream sequence BCAL3494, used during deletion | pGPI<sub>UL-DL</sub> | this study |
| pGPI + BCAM0992 upstream sequence | pGPI-Scel-XCm with ligated upstream sequence BCAM0992, used during deletion | pGPI<sub>UM</sub> | this study |
| pGPI + BCAM0992 upstream and downstream sequence | pGPI-Scel-XCm with ligated upstream and downstream sequence BCAM0992, used during deletion | pGPI<sub>UM-DM</sub> | this study |
| pJH2 + BCAL3494 sequence | Fusion vector with ligated BCAL3494 sequence,<br>used for complementation | pJH2<sub>L3494</sub> | this study |
| pSCrhaB2 + BCAM0992 sequence | Fusion vector with ligated BCAM0992 sequence,<br>used for complementation | pSCrhaB2<sub>M0992</sub> | this study |

### Selection of DNA MTase genes – *in silico*

The REBASE Genome database was used to allocate all known DNA MTase genes in the *B. cenocepacia* J2315 and K56-2 genomes (48). The Artemis Genome Browser and Annotation Tool (Sanger) allowed to visualize the genomic context of these genes (49). NCBI BLAST was used to screen for conservation of the genes within the *Burkholderia* genus using default search parameters (50) (search mode: BLASTn, E cut-off value: < 1E-5).

### Construction of deletion mutants

All primers used for construction and complementation of the deletion mutants are listed in Table S6. The procedure is an adapted allelic replacement approach, using a suicide plasmid with a SceI endonuclease recognition site (51,52). The suicide plasmid, containing DNA fragments of regions flanking the target gene, is integrated into the *B. cenocepacia* genome by homologous recombination. Introducing a second plasmid that carries SceI endonuclease genes into *B. cenocepacia*, results in a lethal genomic strand break. Another homologous recombination event allows the bacteria to repair the break with a 50 % chance of resulting in a gene deletion. Deletion mutants ΔBCAL3494 and ΔBCAM0992 were constructed in both *B. cenocepacia* J2315 and K56-2.

BCAL3494 was deleted together with neighboring gene BCAL3493, as well as BCAL3488 to BCAL3492 (encoding hypothetical proteins). Targeting BCAL3494 alone was not feasible because regions flanking BCAL3494 contain multiple recognition sites for endonucleases used during construction of the deletion mutants, and digestion of these regions would be inevitable (Figure S1).

*E. coli* One Shot PIR2 cells (Thermo Fisher), expressing λ *pir*, were used for transformation, replication, and maintenance of the suicide plasmid during construction of deletion mutants. Thawed cells were immediately exposed to a heat shock transformation procedure, after which they were transferred to SOC medium for recovery. For plasmid selection, the phosphate buffered mineral medium was supplemented with one or more of following antibiotics: trimethoprim (Ludeco; 50 µg/mL for initial screening in *E. coli*, 200 µg/mL when plasmid is introduced in *B. cenocepacia*), chloramphenicol (400 µg/mL), gentamicin (50 µg/mL), kanamycin (50 µg/mL), and ampicillin (200 µg/mL) (Sigma-Aldrich).

### Construction of plasmids for complementation

To ensure that phenotypes were solely caused by the deletion of DNA MTases, deletion mutants were complemented. The primers used for construction of plasmids used for complementation are listed in Table S6. Plasmids pJH2 and pSCrhaB2 were used for complementation of ΔBCAL3494 (*c*ΔBCAL3494) and ΔBCAM0992 (*c*ΔBCAM0992), respectively. The genomic sequences of the DNA MTase genes were PCR-amplified and subsequently cloned into the plasmids. BCAL3494 was amplified including its own regulatory region (approx. 250 nucleotides upstream of the transcription start site) into pJH2, which does not have a promoter associated with its multiple cloning site (53). BCAM0992 does not have its own upstream promoter, therefore it was cloned into pSCrhaB2, which contains a rhamnose-inducible promoter (54). Complemented mutant strains were subjected to the same phenotypic tests as the deletion mutants and wild type *B. cenocepacia*. For strains *c*ΔBCAM0992, the phosphate buffered mineral medium was supplemented with 0.2 % rhamnose.

### Biofilm and clustering experiments

Biofilms were grown in plastic U-shaped 96-well microtiter plates in phosphate buffered medium at 37 °C, starting from 200 µL/well planktonic overnight cultures with an optical density (OD) of 0.05 (590 nm). After 4 h static incubation, all wells were rinsed with physiological saline (PS, 0.9 % NaCl in water), thereby removing all unattached planktonic cells. Wells were re-filled with 200 µL medium and incubated for an additional 20 h. Where appropriate, biofilms were stained with LIVE/DEAD (SYTO9/propidium iodide, Invitrogen) to visualize the bacteria and distinguish live and dead cells (55). Pellicle formation was determined in glass tubes. Cultures were grown statically for 24 h, after which adhering pellicles were stained and quantified with crystal violet (56). Cell clustering, already shown to be correlated with pellicle formation, was determined with flow cytometry (Attune NxT Flow Cytometer, Thermo Fisher) (57). Forward scatter (FCS), a value for particle size, and side scatter (SSC), a value for particle complexity, were measured for each particle present in the bacterial suspension and visualized in scatter plots. After analysis of these graphs, the main cell population was gated (gate ranging from approx. 10^3^ to 10^5^ for both FSC and SSC), and detected events larger and more complex than the gate, were considered clustered (Figure S6).

### Motility experiments

Petri dishes containing phosphate buffered mineral medium with agar concentrations of 0.3 % and 0.5 % were used for assessment of swimming and swarming motility, respectively. 1 µL of cultures with OD 0.1 was spotted on the agar plates. Diameters were measured after 24 h (strain K56-2) or 32 h (strain J2315).

### DNA MTase inhibition with sinefungin

A stock solution of the DNA MTase inhibitor sinefungin (Sigma-Aldrich) was prepared (10 mg/mL) (29), aliquoted, and immediately frozen at −20 °C to prevent degradation. Cells were grown for 24 h in sinefungin-supplemented medium (50 µg/mL) and used as inoculum for an overnight culture, also in sinefungin supplemented medium. This allowed the DNA MTase inhibitor to have an effect during several growth cycles. Then, biofilm formation and motility of sinefungin-treated cells was assessed as described above in medium supplemented with 50 µg/mL sinefungin.

### Genomic DNA extraction

Prior to DNA extraction, planktonic strains were grown overnight in a shaker incubator (100 rpm) at 37 °C. Biofilm cells were grown as described above. Next, cells were harvested and genomic DNA (gDNA) was extracted using the Wizard Genomic DNA Purification Kit (Promega). Quantification was performed with a BioDrop µLITE (BioDrop) spectrophotometer.

### SMRT sequencing

To determine the methylome of *B. cenocepacia*, gDNA extracts were analyzed with Single Molecule Real-Time (SMRT) Sequencing technology. gDNA samples of both wild type and mutant strains were run on a Pacific Biosciences Sequel System (250x coverage) according to the manufacturer’s guidelines. Library preparations were multiplexed as data output of approximately 2 Gb per genome was expected, and a single SMRT Sequel cell provides up to 6 Gb data. Initial data output was processed with SMRT Link software (Pacific Biosciences). Identification of the modified bases and analysis of the methylated motifs was performed with the Base Modification and Motif Analysis application (SMRT Link v6.0, Pacific Biosciences). In depth data analysis was performed with CLC Workbench Genomics (v11.0.1, Qiagen). Differential analysis between wild type and mutants was performed to identify methylation motifs specifically associated with certain DNA MTases. Previously predicted promoter regions and transcription start sites of *B. cenocepacia* were used to determine the methylation profile of regulatory regions (58). Virtual Footprint software (promoter analysis mode, default search parameters) was used to assess similarity of the methylation motifs to known TF binding sites (59).

### qPCR

To evaluate the impact of DNA methylation in promoter regions on gene expression, qPCR was performed on all genes that had a methylated promoter region in wild type *B. cenocepacia*, but an absence of methylation in promoter region in one of the deletion mutants. All hypothetical genes and genes with unknown function, as well as genes with low innate expression level, were excluded from testing. The primers used for qPCR are listed in Table S7. First, all strains were grown to an OD of 0.6 in phosphate buffered medium, after which they were pelleted by centrifugation and frozen at −80 °C. Next, RNA was extracted using the RiboPure – Bacteria extraction kit (Invitrogen), followed by a DNase treatment to remove trace quantities of gDNA. Quantification and measurement of RNA purity of the extracts was performed with a BioDrop µLITE (BioDrop). Subsequently, cDNA was synthesized, using 500 ng RNA per reaction, with a Reverse Transcriptase kit (High-Capacity cDNA RT Kit, Applied Biosystems). Per qPCR reaction, 2 µL template cDNA was mixed with 10 µL GoTaq qPCR Master Mix, 0.6 µL qPCR primer mix (10 µg/mL), and 7.4 µL nuclease-free water according to the GoTaq qPCR Master Mix (Promega) protocol. Samples were run on a CFX96 Real-Time System C1000 Thermal Cycler (Bio-Rad) and output data was processed with Bio-Rad CFX Manager 3.1 software. The baseline threshold was set to a defined 100 RFU. Obtained Cq values were normalized to reference gene *rpo*D (BCAM0918) of which the expression was stable across all samples, differences to wild type were calculated (ΔΔCq) and log-transformed. Volcano plots were used to plot the negative logarithm of statistical p-values against log 2-fold changes (Figure S5).

### Construction of translational eGFP reporter fusions and measurement of eGFP production

Genes with methylated promoter regions that showed a significant upregulation of gene expression in one of both mutant strains, were selected for eGFP experiments. Translational eGFP reporter fusion plasmids were constructed by cloning the regulatory regions of the genes, comprising 60 to 390 nucleotides upstream of the transcription start site, into vector pJH2. The insert is cloned right in front of the eGFP gene and contains an ATG start codon at the 3’-end, in frame with the codon sequence of the gene. All primers used for amplification of the regulatory regions and screening of pJH2 with correct insert length, are listed in Table S8. The plasmids were transferred to *B. cenocepacia* J2315 and K56-2 by triparental mating. Exconjugants were grown on selective plates (LB medium supplemented with 200 µg/mL chloramphenicol and 50 µg/mL gentamicin) and PCR-screened to confirm the presence of the insert. Constructs carrying genes BCAL2415, BCAL2465 and BCAM1362 repeatedly failed to be transformed into *B. cenocepacia* and were not included in further experiments. Fluorescent signals of eGFP production in wild type and mutant strains were measured by flow cytometry (Attune NxT Flow Cytometer, Thermo Fisher) (53).

### Data analysis and statistics

Statistical analysis was performed using SPSS Statistics v. 25 software. All tests and experiments were run in triplicate unless otherwise mentioned. Normality of data was verified with a Shapiro-Wilk test. To check for significant differences between data, normally distributed data was subjected to a T-test or One-way ANOVA test, not normally distributed data to a non-parametric Mann-Whitney U-test. Resulting p-values smaller than 0.05 were reported as statistically significant.

## Acknowledgements

This work was funded by the Special Research Fund of Ghent University (Bijzonder Onderzoeksfonds, BOF, grant number BOFDOC2016001301) and the Swiss National Science Foundation (31003A_169307).

## Supplementary data

**FIGURE S1.**
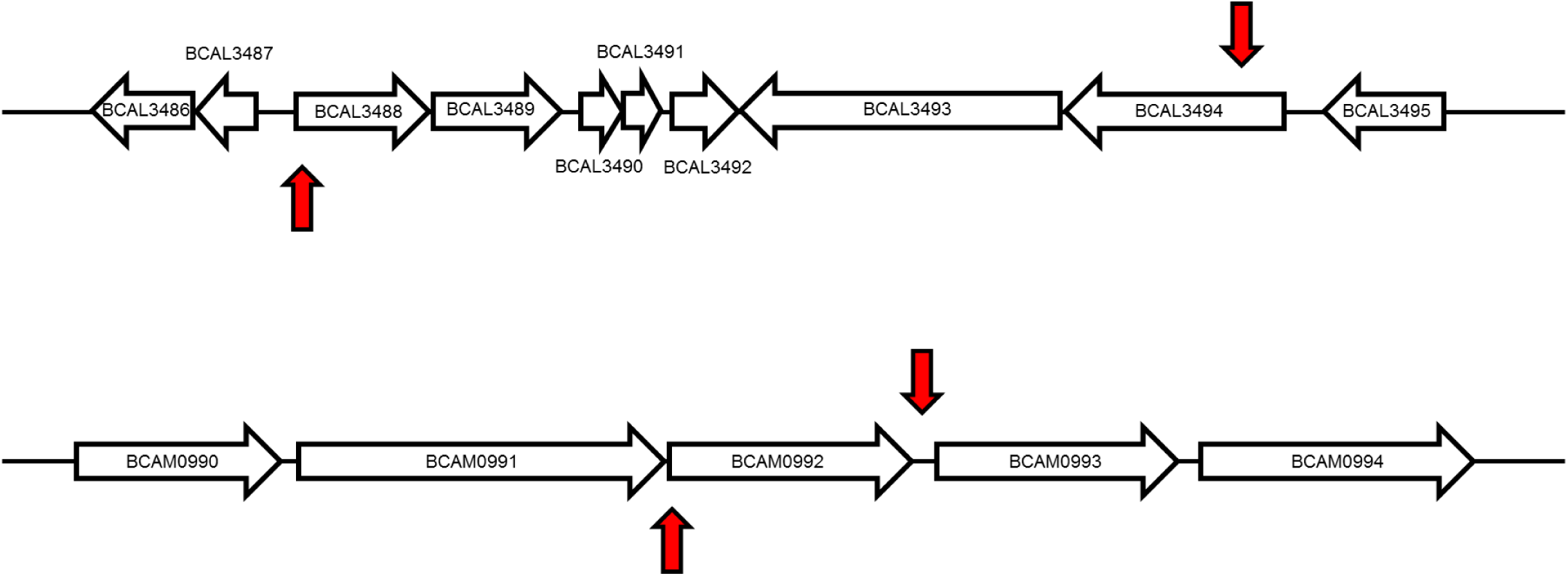
Genome context of deleted DNA MTase genes BCAL3494 (top) and BCAM0992 (bottom). The red arrows indicate the boundaries of the deleted part. For gene BCAL3494, adjacent restriction gene BCAL3493, and hypothetical genes BCAL3488 to BCAL3492 were deleted as well.

**FIGURE S2.**
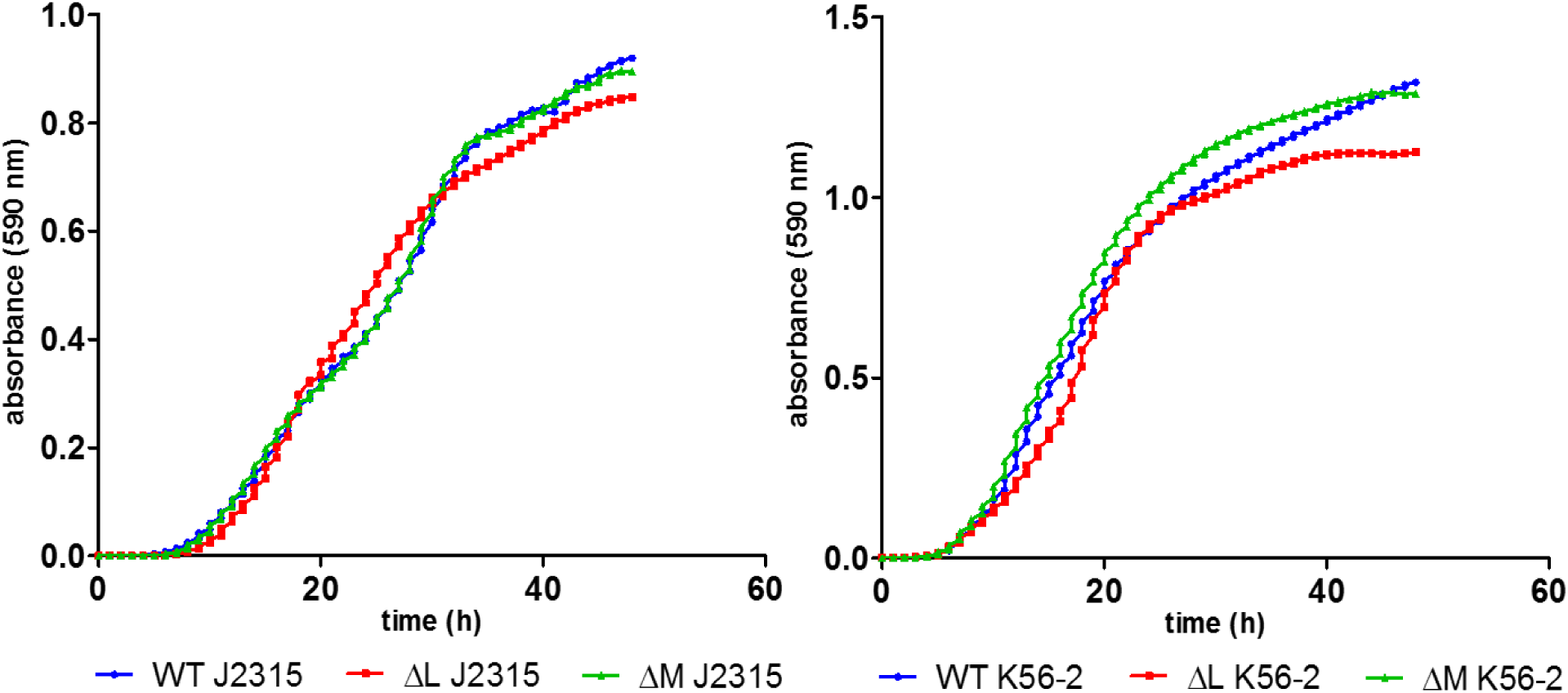
Growth of *B. cenocepacia* J2315 (left) and K56-2 (right) in phosphate buffered minimal medium. (WT: wild type, ΔL: deletion mutant ΔBCAL3494, ΔM: deletion mutant ΔBCAM0992)

**FIGURE S3.**
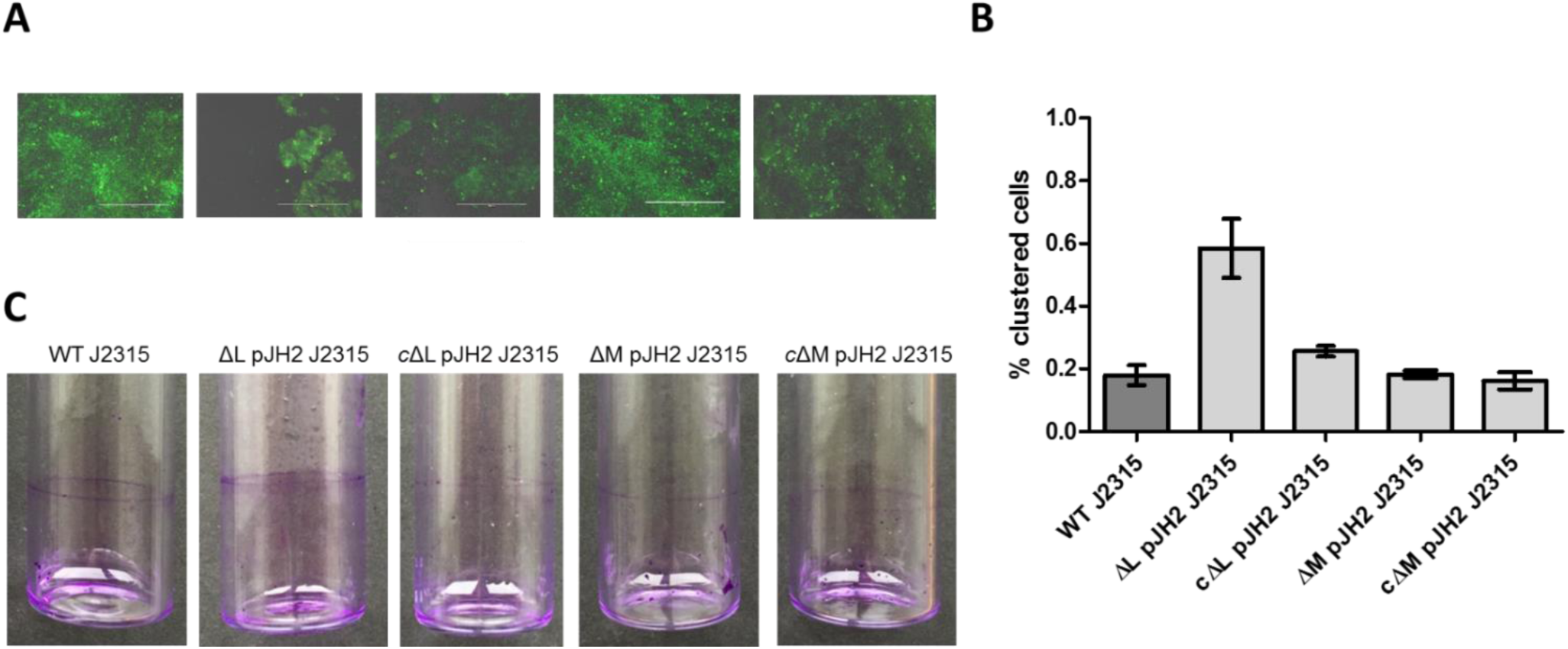
Biofilm formation, cell aggregation, and pellicle formation of complemented DNA MTase mutants in *B. cenocepacia* J2315. (A) Microscopic images of LIVE/DEAD stained biofilms, grown in microtiter plate wells for 24 h. White bar (200 µm) for scale. (B) Clustering of cells in planktonic cultures, analyzed with flow cytometry. (C) Pellicle formation inside glass tubes after 24 h of static incubation, stained with crystal violet. (n=3, error bars represent the Standard Error of the Mean (SEM). WT: wild type, ΔL pJH2 and ΔM pJH2: mutant strains with empty vector pJH2 (vector control), *c*ΔL pJH2 and *c*ΔM pJH2: deletion mutants complemented with genes BCAL3494 and BCAM0992)

**FIGURE S4.**
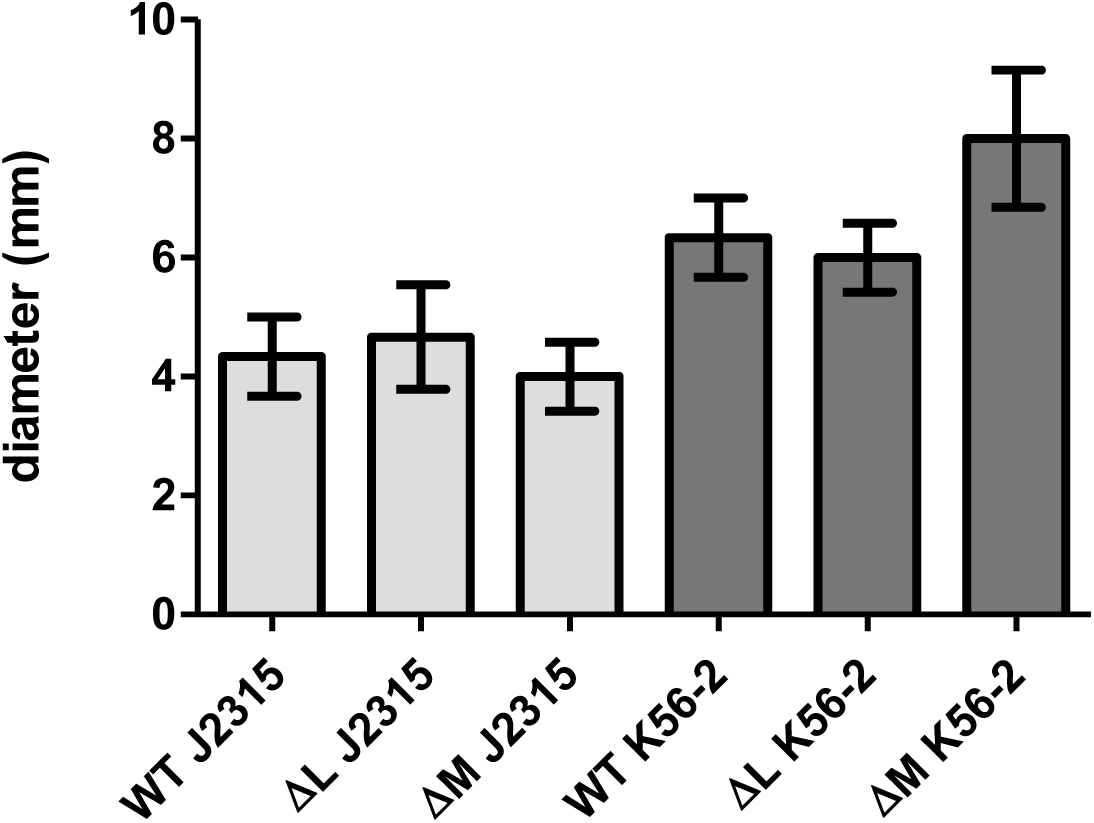
Swarming motility of DNA MTase deletion mutants. Diameters were measured after 24 h. (n=3, error bars represent the SEM. WT: wild type, ΔL: deletion mutant ΔBCAL3494, ΔM: deletion mutant ΔBCAM0992)

**FIGURE S5.**
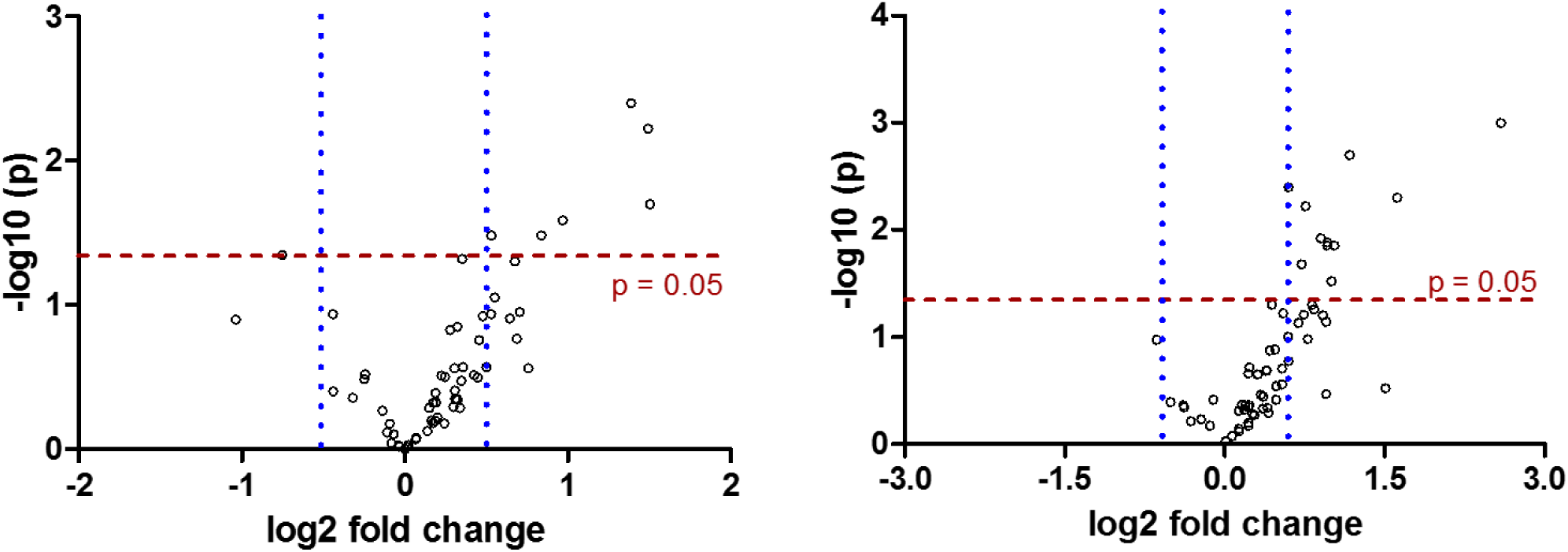
Differential expression (Volcano plots) of all genes with methylated promoter region in J2315 (left) and K56-2 (right) for which expression was quantified using qPCR. Cut-offs were drawn at fold changes −1.5 and 1.5 (blue) and at p-value 0.05 (red). All genes outside of these cut-offs were considered significantly up- or downregulated.

**FIGURE S6.**
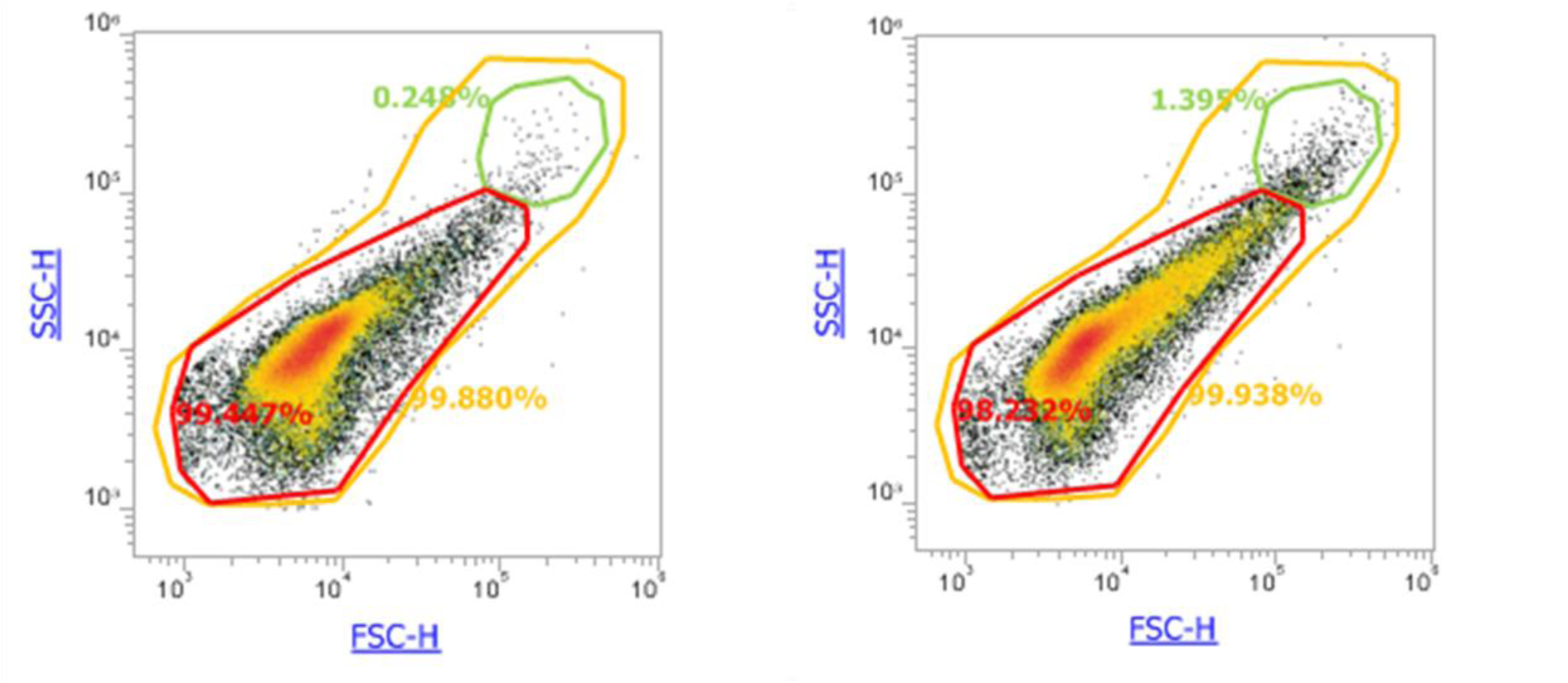
Quantification of the number of clusters in wild type J2315 (left) and ΔBCAL3494 J2315 (right) (SSC: side scatter, FSC: forward scatter, green circle indicates clusters, red circle indicates main population (ranging from approx. 10^3^ to 10^5^ for both FSC and SSC)).

**TABLE S1.** All DNA MTase genes in the *B. cenocepacia* J2315 genome identified by REBASE (* BLASTn search against the genus *Burkholderia*, default screening parameters were used).

| Locus Tag | Name | Location | Predicted type<br>MTase (48) | Widely distributed in<br>genus *Burkholderia* * |
| --- | --- | --- | --- | --- |
| **BCAL3494** | M.BceJI | CHR 1 | Type III | ✓ |
| **BCAM1036** | M.BceJII | CHR 2 |  |  |
| **pBCA072** | M.BceJIII | PLASMID |  |  |
| **BCAM0992** | M.BceJIV | CHR 2 |  | ✓ |
| **BCAL0178** | M.BceJ178P | CHR 1 | Type II |  |
| **BCAL0414** | M.BceJ414P | CHR 1 | Type I |  |

**TABLE S2.**
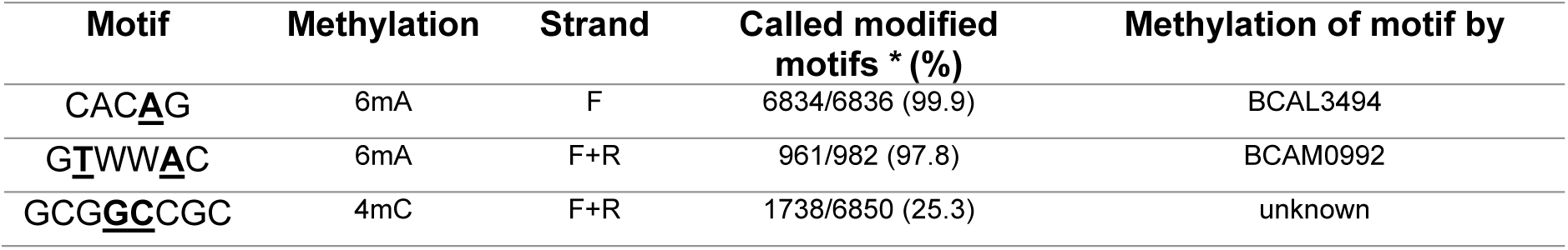
Methylation motifs in *B. cenocepacia* (methylated bases in bold and underlined, 6mA: N_6_-methyl adenine, 4mC: N_4_-methyl cytosine, F: forward strand, R: reverse strand). * wild type strain J2315 percentages.

**TABLE S3.**
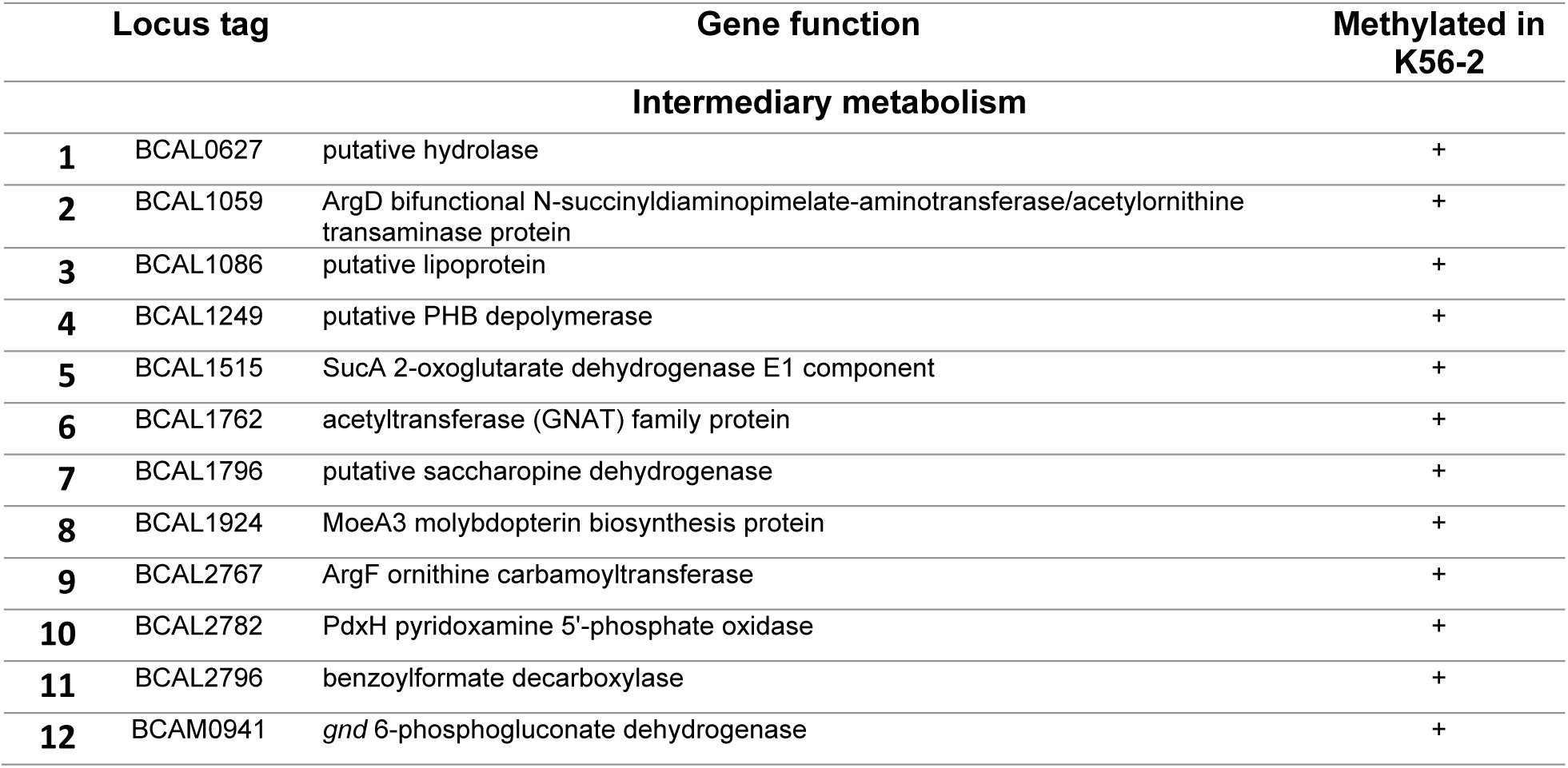
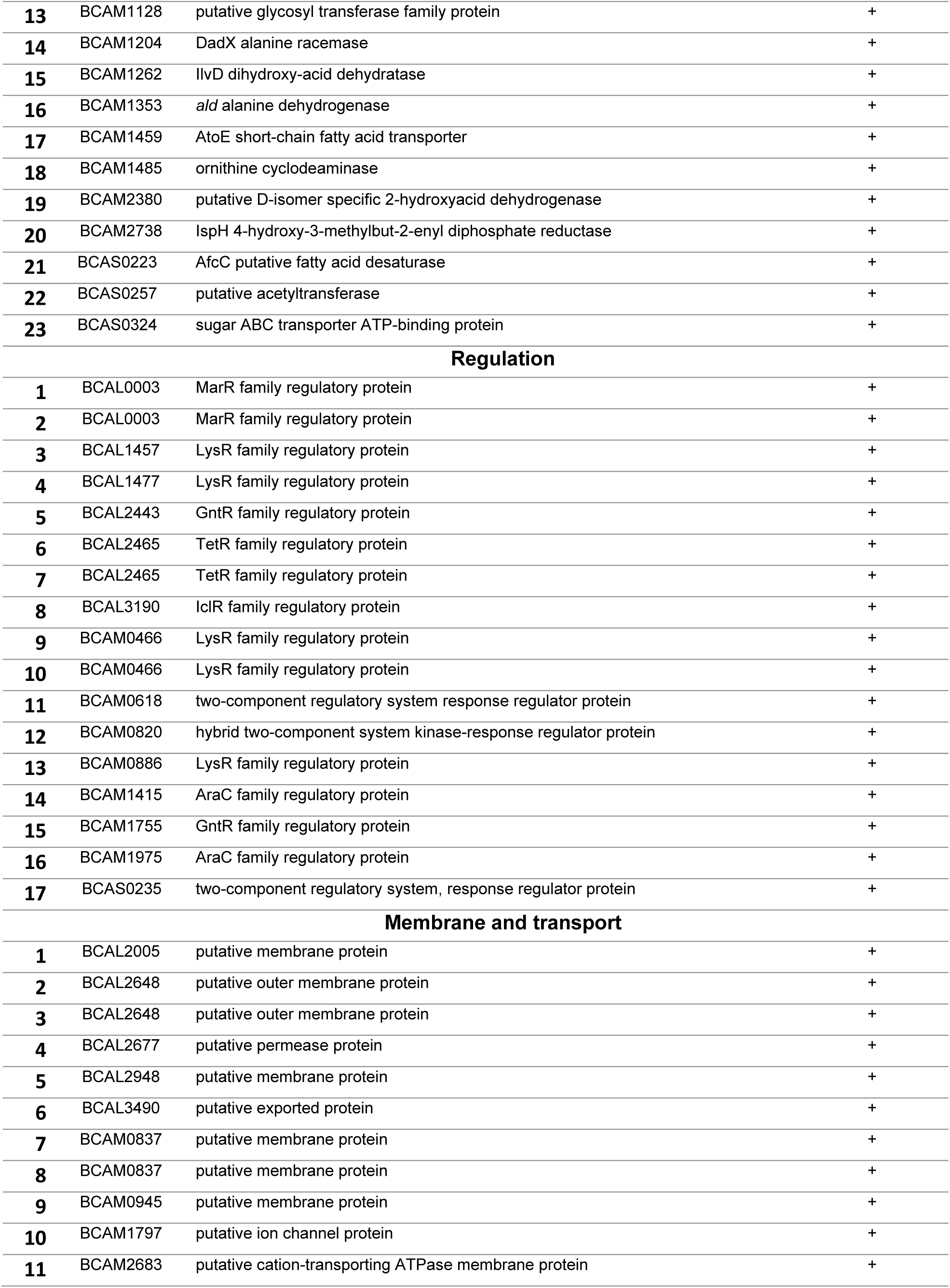
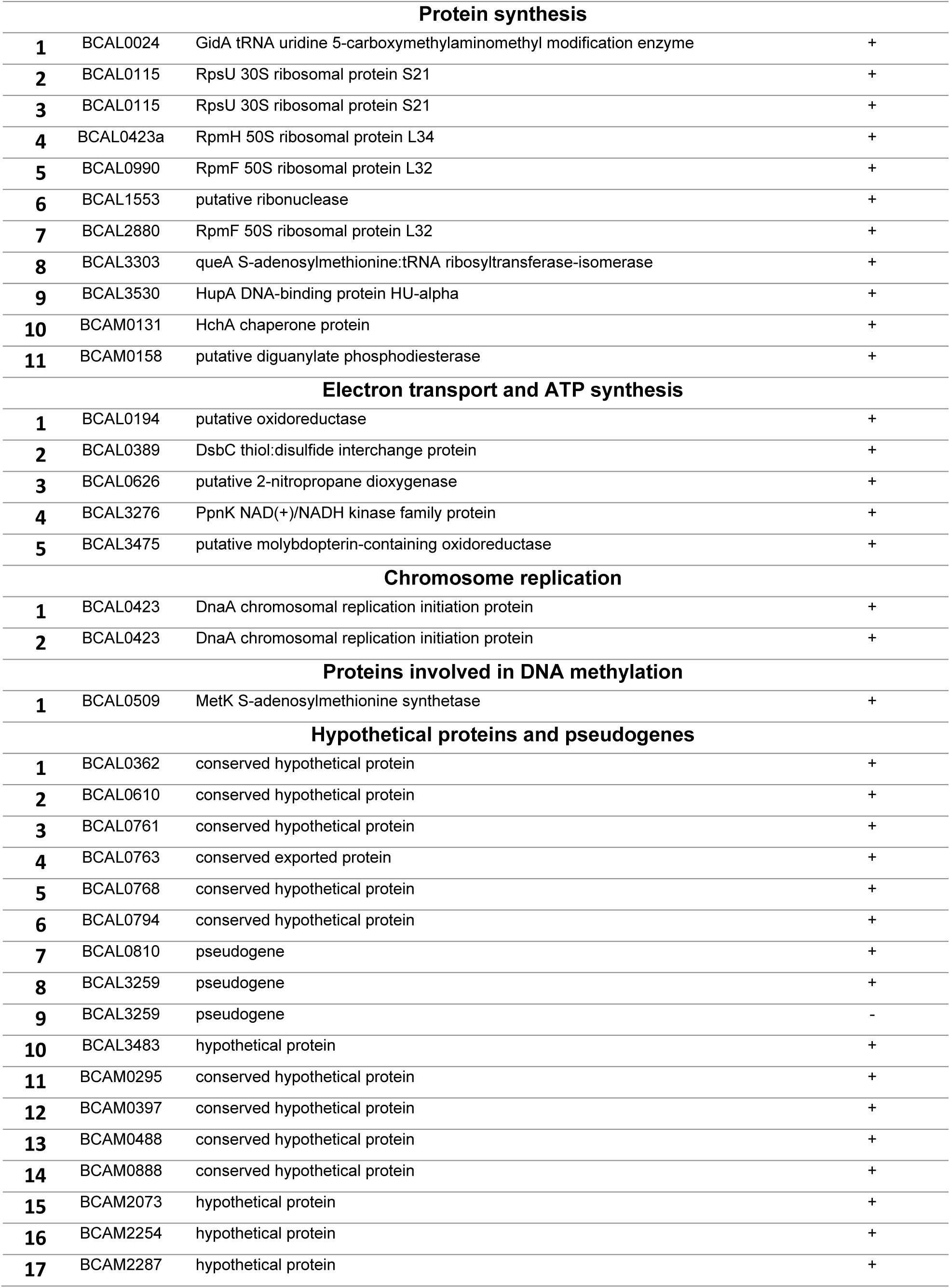
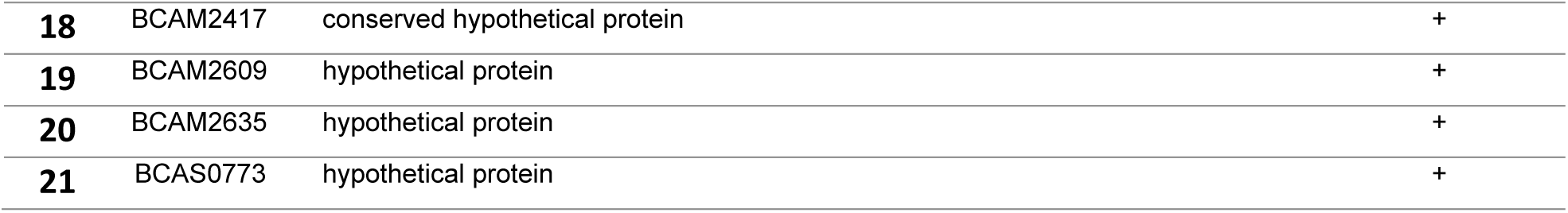
Genes with methylated promoter region (CACAG motif) in J2315 (methylated promoter regions in K56-2 are indicated with ‘+’, non-methylated promoter regions with ‘-’).

| Locus tag | Gene function | Methylated in<br>K56-2 |
| --- | --- | --- |
| **Intermediary metabolism** |  |  |
| **1** BCAL0627 | putative hydrolase | + |
| **2** BCAL1059 | ArgD bifunctional N-succinyl-diaminopimelate-aminotransferase/acetylornithine<br>transaminase protein | + |
| **3** BCAL1086 | putative lipoprotein | + |
| **4** BCAL1249 | putative PHB depolymerase | + |
| **5** BCAL1515 | SucA 2-oxoglutarate dehydrogenase E1 component | + |
| **6** BCAL1762 | acetyltransferase (GNAT) family protein | + |
| **7** BCAL1796 | putative saccharopine dehydrogenase | + |
| **8** BCAL1924 | MoeA3 molybdopterin biosynthesis protein | + |
| **9** BCAL2767 | ArgF ornithine carbamoyltransferase | + |
| **10** BCAL2782 | PdxH pyridoxamine 5'-phosphate oxidase | + |
| **11** BCAL2796 | benzoylformate decarboxylase | + |
| **12** BCAM0941 | *gnd* 6-phosphogluconate dehydrogenase | + |

|  |  |  |  |
| --- | --- | --- | --- |
| 13 | BCAM1128 | putative glycosyl transferase family protein | + |
| 14 | BCAM1204 | DadX alanine racemase | + |
| 15 | BCAM1262 | IlvD dihydroxy-acid dehydratase | + |
| 16 | BCAM1353 | *ald* alanine dehydrogenase | + |
| 17 | BCAM1459 | AtoE short-chain fatty acid transporter | + |
| 18 | BCAM1485 | ornithine cyclodeaminase | + |
| 19 | BCAM2380 | putative D-isomer specific 2-hydroxyacid dehydrogenase | + |
| 20 | BCAM2738 | IspH 4-hydroxy-3-methylbut-2-enyl diphosphate reductase | + |
| 21 | BCAS0223 | AfcC putative fatty acid desaturase | + |
| 22 | BCAS0257 | putative acetyltransferase | + |
| 23 | BCAS0324 | sugar ABC transporter ATP-binding protein | + |
| **Regulation** |  |  |  |
| 1 | BCAL0003 | MarR family regulatory protein | + |
| 2 | BCAL0003 | MarR family regulatory protein | + |
| 3 | BCAL1457 | LysR family regulatory protein | + |
| 4 | BCAL1477 | LysR family regulatory protein | + |
| 5 | BCAL2443 | GntR family regulatory protein | + |
| 6 | BCAL2465 | TetR family regulatory protein | + |
| 7 | BCAL2465 | TetR family regulatory protein | + |
| 8 | BCAL3190 | IclR family regulatory protein | + |
| 9 | BCAM0466 | LysR family regulatory protein | + |
| 10 | BCAM0466 | LysR family regulatory protein | + |
| 11 | BCAM0618 | two-component regulatory system response regulator protein | + |
| 12 | BCAM0820 | hybrid two-component system kinase-response regulator protein | + |
| 13 | BCAM0886 | LysR family regulatory protein | + |
| 14 | BCAM1415 | AraC family regulatory protein | + |
| 15 | BCAM1755 | GntR family regulatory protein | + |
| 16 | BCAM1975 | AraC family regulatory protein | + |
| 17 | BCAS0235 | two-component regulatory system, response regulator protein | + |
| **Membrane and transport** |  |  |  |
| 1 | BCAL2005 | putative membrane protein | + |
| 2 | BCAL2648 | putative outer membrane protein | + |
| 3 | BCAL2648 | putative outer membrane protein | + |
| 4 | BCAL2677 | putative permease protein | + |
| 5 | BCAL2948 | putative membrane protein | + |
| 6 | BCAL3490 | putative exported protein | + |
| 7 | BCAM0837 | putative membrane protein | + |
| 8 | BCAM0837 | putative membrane protein | + |
| 9 | BCAM0945 | putative membrane protein | + |
| 10 | BCAM1797 | putative ion channel protein | + |
| 11 | BCAM2683 | putative cation-transporting ATPase membrane protein | + |

| **Protein synthesis** |  |  |  |
| --- | --- | --- | --- |
| **1** | BCAL0024 | GidA tRNA uridine 5-carboxymethylaminomethyl modification enzyme | + |
| **2** | BCAL0115 | RpsU 30S ribosomal protein S21 | + |
| **3** | BCAL0115 | RpsU 30S ribosomal protein S21 | + |
| **4** | BCAL0423a | RpmH 50S ribosomal protein L34 | + |
| **5** | BCAL0990 | RpmF 50S ribosomal protein L32 | + |
| **6** | BCAL1553 | putative ribonuclease | + |
| **7** | BCAL2880 | RpmF 50S ribosomal protein L32 | + |
| **8** | BCAL3303 | queA S-adenosylmethionine:tRNA ribosyltransferase-isomerase | + |
| **9** | BCAL3530 | HupA DNA-binding protein HU-alpha | + |
| **10** | BCAM0131 | HchA chaperone protein | + |
| **11** | BCAM0158 | putative diguanylate phosphodiesterase | + |
| **Electron transport and ATP synthesis** |  |  |  |
| **1** | BCAL0194 | putative oxidoreductase | + |
| **2** | BCAL0389 | DsbC thiol:disulfide interchange protein | + |
| **3** | BCAL0626 | putative 2-nitropropane dioxygenase | + |
| **4** | BCAL3276 | PpnK NAD(+)/NADH kinase family protein | + |
| **5** | BCAL3475 | putative molybdopterin-containing oxidoreductase | + |
| **Chromosome replication** |  |  |  |
| **1** | BCAL0423 | DnaA chromosomal replication initiation protein | + |
| **2** | BCAL0423 | DnaA chromosomal replication initiation protein | + |
| **Proteins involved in DNA methylation** |  |  |  |
| **1** | BCAL0509 | MetK S-adenosylmethionine synthetase | + |
| **Hypothetical proteins and pseudogenes** |  |  |  |
| **1** | BCAL0362 | conserved hypothetical protein | + |
| **2** | BCAL0610 | conserved hypothetical protein | + |
| **3** | BCAL0761 | conserved hypothetical protein | + |
| **4** | BCAL0763 | conserved exported protein | + |
| **5** | BCAL0768 | conserved hypothetical protein | + |
| **6** | BCAL0794 | conserved hypothetical protein | + |
| **7** | BCAL0810 | pseudogene | + |
| **8** | BCAL3259 | pseudogene | + |
| **9** | BCAL3259 | pseudogene | - |
| **10** | BCAL3483 | hypothetical protein | + |
| **11** | BCAM0295 | conserved hypothetical protein | + |
| **12** | BCAM0397 | conserved hypothetical protein | + |
| **13** | BCAM0488 | conserved hypothetical protein | + |
| **14** | BCAM0888 | conserved hypothetical protein | + |
| **15** | BCAM2073 | hypothetical protein | + |
| **16** | BCAM2254 | hypothetical protein | + |
| **17** | BCAM2287 | hypothetical protein | + |
| **18** | BCAM2417 | conserved hypothetical protein | + |
| **19** | BCAM2609 | hypothetical protein | + |
| **20** | BCAM2635 | hypothetical protein | + |
| **21** | BCAS0773 | hypothetical protein | + |

**TABLE S4.** Genes with methylated promoter region (GTWWAC motif) in J2315 (methylated promoter regions in K56-2 are indicated with ‘+’, non-methylated promoter regions with ‘-’).

| Locus tag |  | Gene function | Methylated in K56-2 |
| --- | --- | --- | --- |
| **Intermediary metabolism** |  |  |  |
| **1** | BCAL0064 | AcoD acetaldehyde dehydrogenase | + |
| **2** | BCAL0162 | GmhA phosphoheptose isomerase | + |
| **3** | BCAL0508 | LpxL lipid A biosynthesis myristoyl acyltransferase | + |
| **4** | BCAL0709 | LipB lipoate-protein ligase B | + |
| **5** | BCAL0995 | AcpP acyl carrier protein | + |
| **6** | BCAL1290 | undecaprenyl pyrophosphate phosphatase | + |
| **7** | BCAL1467 | AroC chorismate synthase | + |
| **8** | BCAL1478 | putative hydrolase | + |
| **9** | BCAL1556 | RpiA ribose-5-phosphate isomerase A | + |
| **10** | BCAL1938 | family C40 cysteine peptidase | + |
| **11** | BCAL2406 | WabR putative glycosyltransferase | + |
| **12** | BCAL2419 | glycosyl hydrolases family protein | + |
| **13** | BCAL2701 | ArgD acetylornithine transaminase protein | + |
| **14** | BCAL2875 | AcpP acyl carrier protein | + |
| **15** | BCAL2942 | CysM cysteine synthase B | + |
| **16** | BCAL3153 | putative lipoprotein | + |
| **17** | BCAL3179 | LdhA putative D-lactate dehydrogenase | + |
| **18** | BCAM0013 | putative acetyltransferase | + |
| **19** | BCAM1679 | putative lysylphosphatidylglycerol synthetase | + |
| **20** | BCAM1761 | putative lipoprotein | + |
| **21** | BCAM2511 | GarD putative D-galactarate dehydratase | + |
| **22** | BCAM2737 | putative glycosyl transferase | + |
| **23** | BCAS0065 | putative glutathione S-transferase | + |
| **Membrane and transport** |  |  |  |
| **1** | BCAL0121 | AqpZ aquaporin Z | + |
| **2** | BCAL0126 | MotA chemotaxis protein | + |
| **3** | BCAL0824 | putative membrane protein | + |
| **4** | BCAL1525 | *flp* type pilus subunit | + |
| **5** | BCAL1808 | putative membrane protein | + |
| **6** | BCAL2301 | putative exported protein | + |
| **7** | BCAL2370 | putative membrane protein | + |
| **8** | BCAM0837 | putative membrane protein | + |
| **9** | BCAM0885 | putative membrane protein | + |
| **10** | BCAM0988 | putative exported protein | + |
| **11** | BCAM1669 | putative exported protein | + |
| **12** | BCAM1726 | putative exported protein | + |
| **13** | BCAM1743 | periplasmic solute-binding protein | + |
| **14** | BCAM1830 | putative exported protein | + |
| **15** | BCAM2555 | putative exported protein | + |
| **16** | BCAM2827 | putative exported protein | + |
| **17** | BCAM2828 | putative membrane protein | + |
| **Regulation** |  |  |  |
| **1** | BCAL0054 | MerR family regulatory protein | + |
| **2** | BCAL0444 | GntR family regulatory protein | + |
| **3** | BCAL0497 | two-component regulatory system, sensor kinase protein | + |
| **4** | BCAL2606 | two-component regulatory system, response regulator protein | + |
| **5** | BCAM0076 | TetR family regulatory protein | + |
| **6** | BCAM0085 | TetR family regulatory protein | + |
| **7** | BCAM0483 | ADA-like AraC family regulatory protein | + |
| **8** | BCAM2589 | IclR family regulatory protein | + |
| **9** | BCAS0007 | TetR family regulatory protein | + |
| **10** | BCAS0258 | GntR family regulatory protein | + |
| **Electron transport and ATP synthesis** |  |  |  |
| **1** | BCAL0080 | putative cytochrome | + |
| **2** | BCAL2145 | NADH-ubiquinone oxidoreductase subunit | + |
| **3** | BCAL2415 | PurT phosphoribosylglycinamide formyltransferase 2 | + |
| **4** | BCAL3187 | putative oxidoreductase | + |
| **5** | BCAL3362 | putative oxidoreductase | + |
| **6** | BCAL3432 | cytochrome c assembly protein | + |
| **DNA transposition** |  |  |  |
| **1** | BCAL2216 | putative transposase | + |
| **2** | BCAL3238 | putative transposase | + |
| **3** | BCAL3295 | putative transposase | + |
| **Protein synthesis** |  |  |  |
| **1** | BCAL1856 | RimO ribosomal protein S12 methylthiotransferase | + |
| **2** | BCALr3484 | tRNA-Val | + |
| **Chromosome replication** |  |  |  |
| **1** | BCAL0079 | *rep* ATP-dependent DNA helicase | + |
| **Proteins involved in DNA methylation** |  |  |  |
| **1** | BCAL0747 | putative methyltransferase | + |
| **Other proteins** |  |  |  |
| 1 | BCAL0825 | UvrA excinuclease ABC subunit A | + |
| 2 | BCAL0849 | subfamily M48B metallopeptidase | + |
| 3 | BCAL2643 | SodC superoxide dismutase | + |
| 4 | BCAL2749 | putative diguanylate phosphodiesterase | + |
| 5 | BCAL2940 | putative histone deacetylase-family protein | + |
| 6 | BCAM1362 | putative penicillin-binding protein | + |
| 7 | BCAS0010 | putative activator of osmoprotectant transporter | + |
| 8 | BCAS0293 | AidA nematocidal protein | + |
| **Hypothetical proteins and pseudogenes** |  |  |  |
| 1 | BCAL0362 | conserved hypothetical protein | + |
| 2 | BCAL0434 | putative exported protein | + |
| 3 | BCAL1442 | conserved hypothetical protein | + |
| 4 | BCAL2175 | conserved hypothetical protein | + |
| 5 | BCAL3193 | conserved hypothetical protein | + |
| 6 | BCAM0008 | conserved hypothetical protein | + |
| 7 | BCAM1874-2 | pseudo | + |
| 8 | BCAM2207 | conserved hypothetical protein | + |
| 9 | BCAM2289 | conserved hypothetical protein | + |

**TABLE S5.**
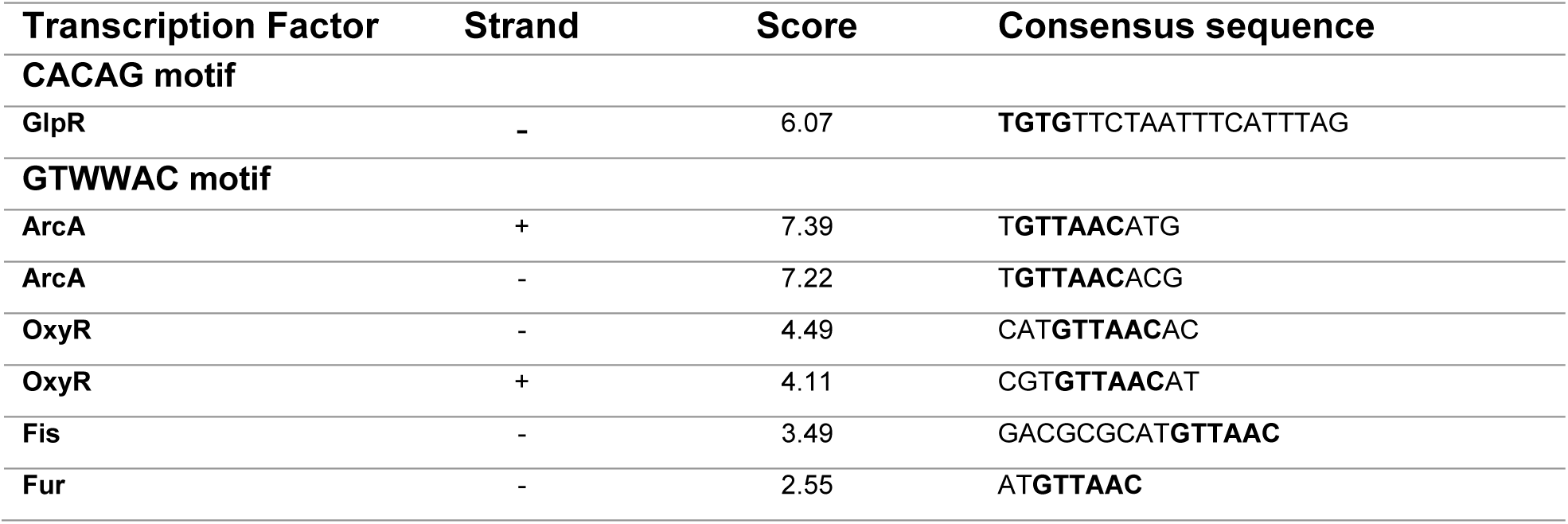
List of TFs that bind to methylation motifs CACAG and GTWWAC, predicted by Virtual Footprint. Bold sequences represent methylation motifs. (Consensus sequence based on TF binding in *E. coli* K12)

**TABLE S6.**
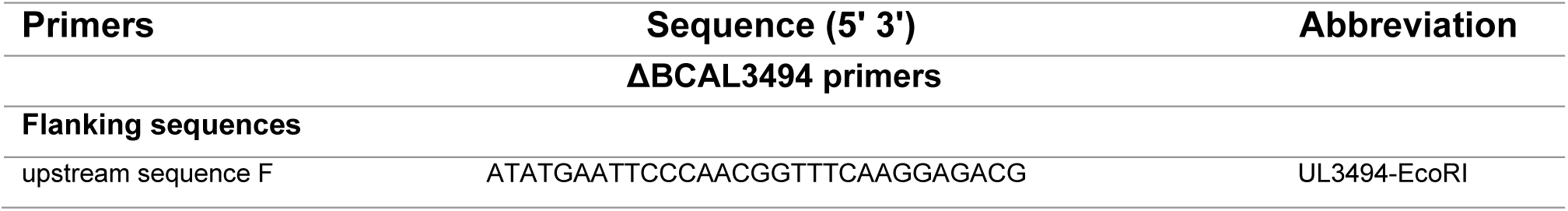
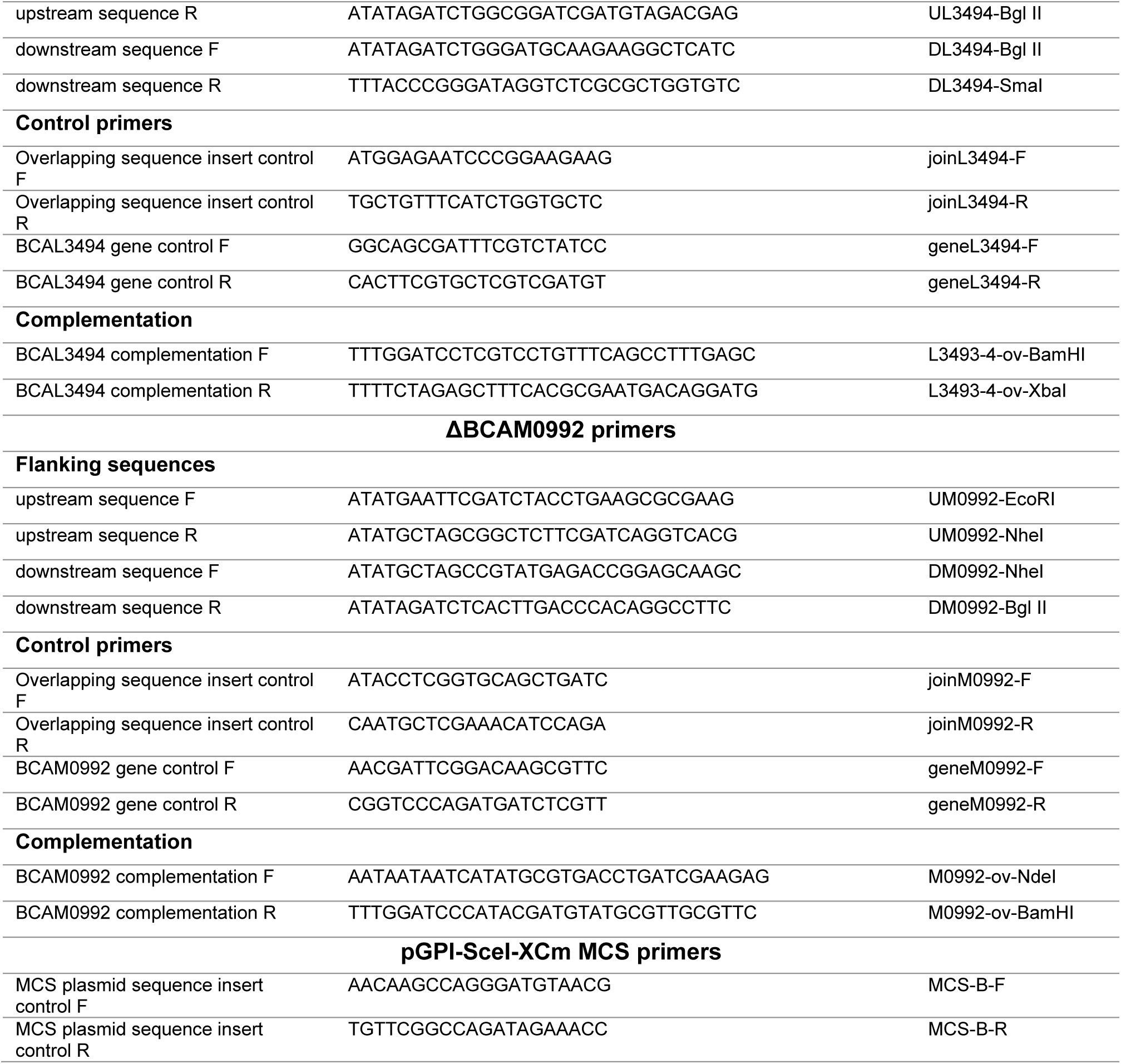
Overview of all primers used for construction and complementation of gene deletion mutants.

**TABLE S7.**
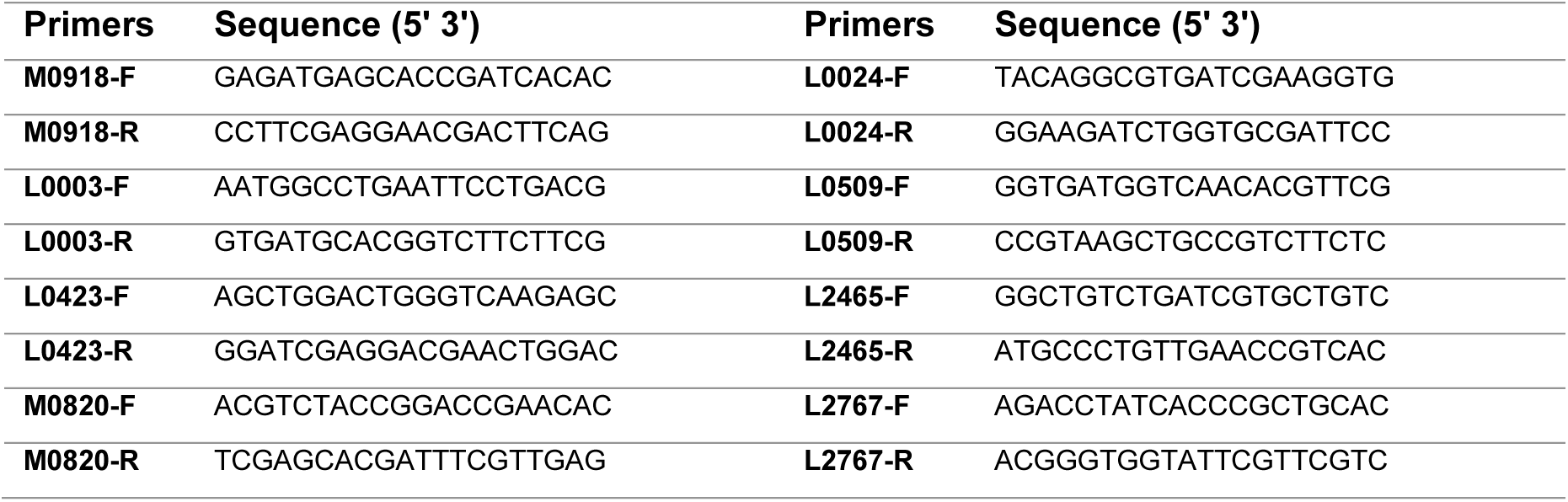
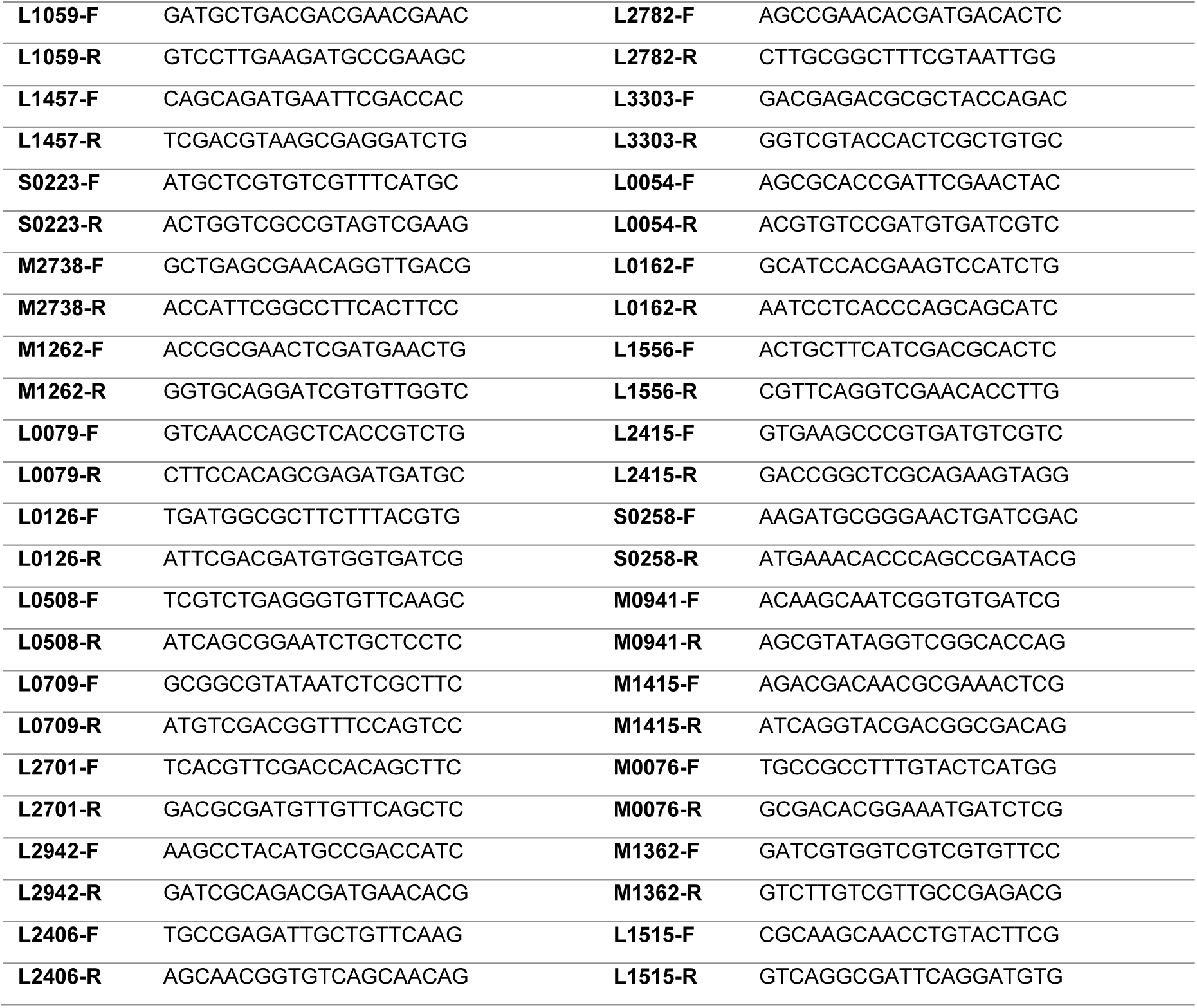
Primers used in qPCR experiments.

**TABLE S8.**
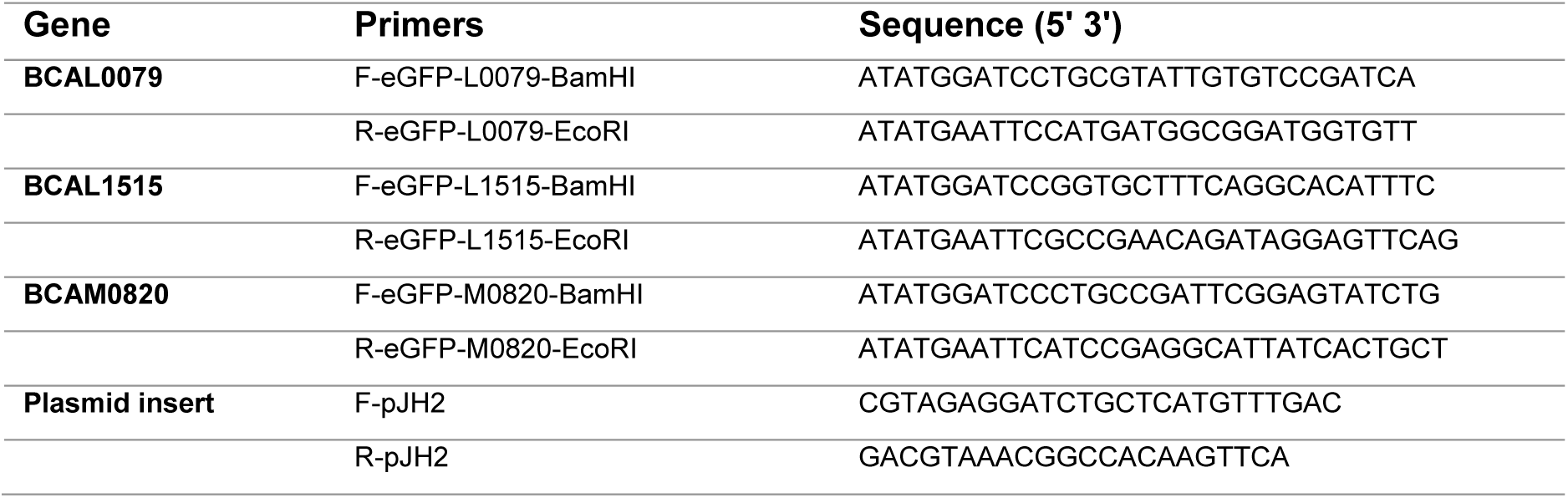
List of primers used for construction of translational eGFP reporter fusion plasmids.

